# Intracellular BAPTA directly inhibits PFKFB3, thereby impeding mTORC1-driven Mcl-1 translation and killing Mcl-1-addicted cancer cells

**DOI:** 10.1101/2022.10.31.512457

**Authors:** Flore Sneyers, Martijn Kerkhofs, Kirsten Welkenhuyzen, Femke Speelman-Rooms, Ahmed Shemy, Arnout Voet, Guy Eelen, Mieke Dewerchin, Stephen W. Tait, Bart Ghesquière, Martin D. Bootman, Geert Bultynck

**Affiliations:** KU Leuven, Laboratory of Molecular and Cellular Signaling, Department Cellular and Molecular Medicine, Campus Gasthuisberg O&N I box 802, Herestraat 49, 3000, Leuven, Belgium; KU Leuven, Laboratory for Biomolecular Modelling and Design, Department of Chemistry, Celestijnenlaan 200G, 3001 Heverlee, Belgium; KU Leuven, Laboratory of Angiogenesis and Vascular Metabolism, Department of Oncology, Leuven Cancer Institute, Campus Gasthuisberg O&N4, Herestraat 49 box 912, Leuven, Belgium; VIB – KU Leuven, Center for Cancer Biology, Laboratory of Angiogenesis and Vascular Metabolism, Campus Gasthuisberg O&N4, Herestraat 49 box 912, 3000 Leuven, Belgium; University of Glasgow, Cancer Research UK Beatson Institute, Institute of Cancer Sciences, Glasgow, UK; VIB – KU Leuven Center for Cancer Biology, Metabolomics Expertise Center, Campus Gasthuisberg O&N 4 box 912, Herestraat 49, 3000 Leuven, Belgium; The Open University, School of Life, Health and Chemical Sciences, Faculty of Science, Technology, Engineering and Mathematics, Walton Hall, Milton Keynes, MK7 6AA, United Kingdom

## Abstract

Intracellular Ca^2+^ signals control several physiological and pathophysiological processes. The main tool to chelate intracellular Ca^2+^ is intracellular BAPTA (BAPTA_i_), usually introduced into cells as a membrane-permeant acetoxymethyl ester (BAPTA-AM). We previously demonstrated that BAPTA_i_ enhanced apoptosis induced by venetoclax, a Bcl-2 antagonist, in diffuse large B-cell lymphoma (DLBCL). These findings implied a novel interplay between intracellular Ca^2+^ signaling and anti-apoptotic Bcl-2 function. Hence, we set out to identify the underlying mechanisms by which BAPTA_i_ enhances cell death in B-cell cancers. In this study, we observed that BAPTA_i_ alone induced apoptosis in lymphoma cell models that were highly sensitive to S63845, an Mcl-1 antagonist. BAPTA_i_ provoked a rapid decline in Mcl-1 protein levels by inhibiting mTORC1-driven *MCL-1* translation. Overexpression of nondegradable Mcl-1 rescued BAPTA_i_-induced cell death. We further examined how BAPTA_i_ diminished mTORC1 activity and found that BAPTA_i_ impaired glycolysis by directly inhibiting 6-phosphofructo-2-kinase/fructose-2,6-bisphosphatase 3 (PFKFB3) activity, an up to now unkown effect of BAPTA_i_. All aforementioned effects of BAPTA_i_ were also elicited by a BAPTA_i_ analog with low affinity for Ca^2+^. Thus, our work reveals PFKFB3 inhibition as an unappreciated Ca^2+^-independent mechanism by which BAPTA_i_ impairs cellular metabolism and ultimately the survival of Mcl-1-dependent cancer cells. Our work has two important implications. First, direct inhibition of PFKFB3 emerged as a key regulator of mTORC1 activity and a promising target in the treatment of Mcl-1-dependent cancers. Second, cellular effects caused by BAPTA_i_ are not necessarily related to Ca^2+^ signaling. Our data support the need for a reassessment of the role of Ca^2+^ in cellular processes when findings were based on the use of BAPTA_i_.

## Introduction

Intracellular Ca^2+^ is a ubiquitous second messenger that controls cellular processes ranging from fertilization and cell division to cell death(1),(2).

Cancer cells attenuate intracellular Ca^2+^ signaling, thereby promoting tumorigenesis and conferring resistance to chemotherapeutic regimens(3),(4). Several tumor suppressor proteins and (proto)-oncogenes, including anti-apoptotic members of the Bcl-2 family, directly modulate Ca^2+^ fluxes to enhance survival(5)–(8).

Moreover, anti-apoptotic Bcl-2 family members, including Bcl-2, Bcl-XL and Mcl-1, inhibit the mitochondrial apoptotic pathway(9). Increased abundance of anti-apoptotic Bcl-2, the founding member of the family, is found in several B-cell malignancies, including diffuse large B-cell lymphoma (DLBCL), thereby counteracting pro-apoptotic signaling induced by oncogenic stress(10). This “primed to death” status represents an Achilles’ heel in cancers that can be exploited by BH3-mimetic drugs (such as venetoclax), which antagonize anti-apoptotic Bcl-2 proteins. However, venetoclax resistance in cancer cells has already emerged due to acquired mutations in Bcl-2 and upregulation of Bcl-XL or Mcl-1(11). Hence, selective BH3-mimetic antagonists have been developed for Bcl-XL (including A-1331852, A-1155463 and WEHI-539) and Mcl-1 (S63845), thereby driving cell death in Bcl-XL- and Mcl-1-dependent cancer cells(12),(13).

We previously demonstrated that intracellular BAPTA (BAPTA_i_), a widely used, high-affinity chelator of cytosolic Ca^2+^ loaded into cells as BAPTA-acetoxymethyl ester (BAPTA-AM), enhanced venetoclax-induced cell death in OCI-LY-1 and SU-DHL-4 cells, two DLBCL cell models(14). Extracellularly added BAPTA-AM can pass across the plasma membrane and thus enter the cell, where it is hydrolyzed into the free acid form of BAPTA that chelates cytosolic Ca^2+^. As such, BAPTA is trapped in the cells. BAPTA-AM is typically applied in the extracellular environment at about 10 μM. However, due to the accumulation of the hydrolyzed free acid form, which is cell membrane-impermeant, intracellular BAPTA concentrations that can reach 1 to 2 mM (15).

Given the fast Ca^2+^-chelating properties enabling BAPTA_i_ to buffer Ca^2+^ in close proximity to Ca^2+^channels and microdomains, our results suggested an unprecedented interplay between constitutive Ca^2+^ signaling and the anti-apoptotic function of Bcl-2 proteins in B-cell cancer cells. However, intracellularly loaded BAPTA, and its derivatives, may also interfere with cell physiological processes independently of its Ca^2+^-chelating properties by directly targeting proteins such as the Na^+^/K^+^ATPase(15),(16).

Despite the emerging evidence for off-target effects, BAPTA_i_ continues to be widely used without adequate controls(16). Indeed, thousands of publications have used such chemical Ca^2+^ buffers to delineate the involvement of Ca^2+^ signals in specific cellular processes(17)–(20). Moreover, since BAPTA_i_ can theoretically buffer microscopic ‘local Ca^2+^ signals’ originating in close proximity to Ca^2+^channels, BAPTA_i_ has routinely been used to determine the involvement of Ca^2+^ in a cellular process without actually visualizing the Ca^2+^ signals. A significant confounding issue is that by directly affecting targets independently of its Ca^2+^-chelating properties, the cellular actions of BAPTA_i_ and related compounds may not be limited to Ca^2+^ buffering(21),(22). This urges an in-depth investigation into this unanticipated property of chemical Ca^2+^ buffers(16). Understanding the mechanism by which BAPTA_i_ affects B-cell cancer cell survival and augments their sensitivity to Bcl-2 inhibitors is not only important for developing improved anticancer strategies but also key to unraveling which processes underlie this fascinating interplay.

Here, we aimed to decipher the molecular mechanisms by which BAPTA_i_ impacted cancer cell survival. We found that BAPTA_i_ by itself could evoke cell death in Mcl-1-dependent cancer cells, but not in those cells not addicted to Mcl-1. To distinguish between Ca^2+^-dependent versus Ca^2+^-independent effects of BAPTA_i_, we used a variety of BAPTA derivatives (Figure 1), including tetrafluoro-BAPTA_i_ (TF-BAPTA_i_), whose affinity for Ca^2+^ is about 400-fold lower (K_D_ ~65 μM) than that of BAPTA (K_D_ ~160 nM). As such, the affinity of TF-BAPTA_i_ for Ca^2+^ is about 500 to 600-fold than the basal cytosolic Ca^2+^ concentration, which is about 100 nM. However, similarly to BAPTA_i_, TF-BAPTA_i_ was equally potent in inducing the death of Mcl-1-addicted cancer cells. Using a broad array of approaches, we revealed a novel Ca^2+^-independent, inhibitory effect of BAPTA_i_ on glycolysis, thereby rapidly suppressing mTORC1 activity and thus abrogating mTORC1-driven processes such as Mcl-1 translation. Due to its short-lived nature, Mcl-1 protein levels rapidly decline in BAPTA_i_-loaded cells. In cancer cells, or genetically engineered cells that are dependent on Mcl-1 for cell survival, BAPTA_i_ (and TF-BAPTA_i_) therefore provokes significant cell death, irrespective of intracellular Ca^2+^ buffering. We further identified the presumed BAPTA target, demonstrating that BAPTA directly inhibited 6-phosphofructo-2-kinase/fructose-2,6-bisphosphatase-3 (PFKFB3). In addition, direct PFKFB3 inhibition using AZ67 mimicked the inhibitory effect of BAPTA_i_ on mTORC1 activity and Mcl-1 protein levels. Overall, our work sheds novel light on the cellular effects and direct targets of BAPTA_i_, revealing a novel link between PFKFB3, mTORC1 and Mcl-1 irrespective of Ca^2+^ chelation.

**Figure 1:**
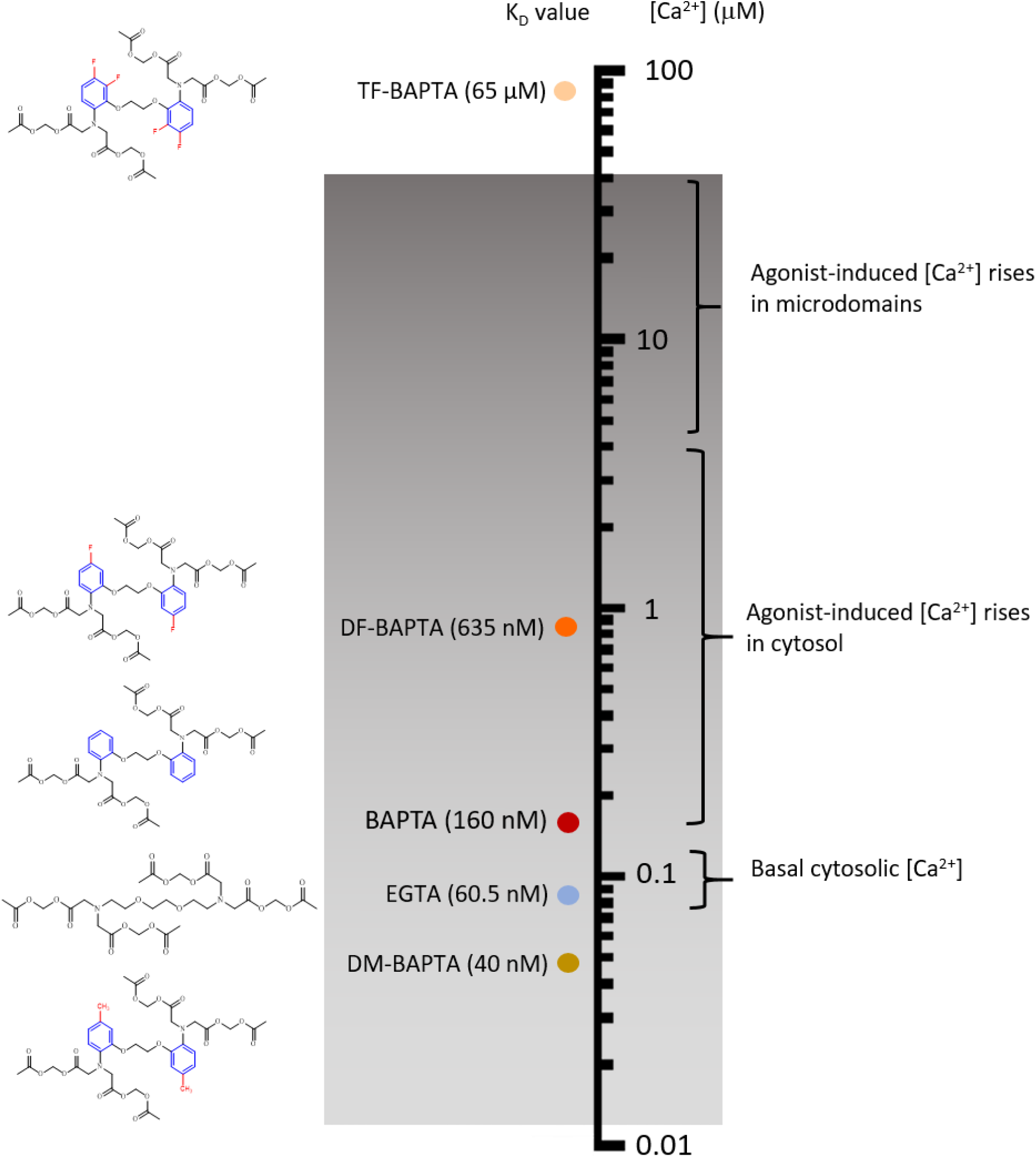
Ca^2+^ chelating compounds and their corresponding K_D_ values in relation to intracellular Ca^2+^ concentrations. Schematic overview of the chemical structures and K_D_ values for Ca^2+^ binding for BAPTA, its derivatives and EGTA in relation to cytosolic Ca^2+^ concentrations in cells. The affinity of TF-BAPTA is about 500 to 600-fold lower than the basal cytosolic Ca^2+^ concentrations.

## Results

### 1. BAPTA_i_ induces apoptosis in B-cell lymphoma cell lines

To explore whether DLBCL cell models could be addicted to Ca^2+^ signaling for their survival, we applied 10 μM BAPTA-AM to the extracellular environment to load the cells with BAPTA_i_ and determined the proportion of annexin V-FITC/7-AAD-stained cells using flow cytometry, as a measure of cell survival. The percentage of living cells was quantified by determining the proportion of AnnexinV/7-AAD-negative cells at different time points and benchmarked against vehicle treatments for each time point. We found that OCI-LY-1 and SU-DHL-4 cells displayed different kinetics and extents of apoptosis (AnnexinV-positive cells) in response to BAPTA_i_ (Figure 2A and 2C for typical cytometry analysis for 6-hour-treated cells; 2B and 2D for results from replicate analysis for different time points). In comparison to vehicle treatment, 6 hours of BAPTA_i_ loading in OCI-LY-1 cells evoked an increase in the proportion of AnnexinV-positive cells, indicative of apoptotic cell death (Figure 2A). In contrast, in SU-DHL-4 cells, 6 hours of BAPTA_i_ loading did not evoke any additional AnnexinV-positive cells in comparison to vehicle treatment (Figure 2D). Such analyses were performed for different BAPTA_i_ loading times. OCI-LY-1 cells exposed to BAPTA_i_ displayed rapid and extensive apoptotic cell death that steadily increased over a time course of 8 hours, a feature not observed in vehicle-treated conditions (Figure 2B). In contrast, SU-DHL-4 cells exposed to BAPTA_i_ did not display any significant apoptosis in a time period of 8 hours (Figure 2E). The apoptotic fraction was determined for each time point (i.e. percentage of living vehicle-treated cells - percentage of living BAPTA_i_-loaded cells) in both OCI-LY-1 and SU-DHL-4 cells (Figure 2E).

**Figure 2:**
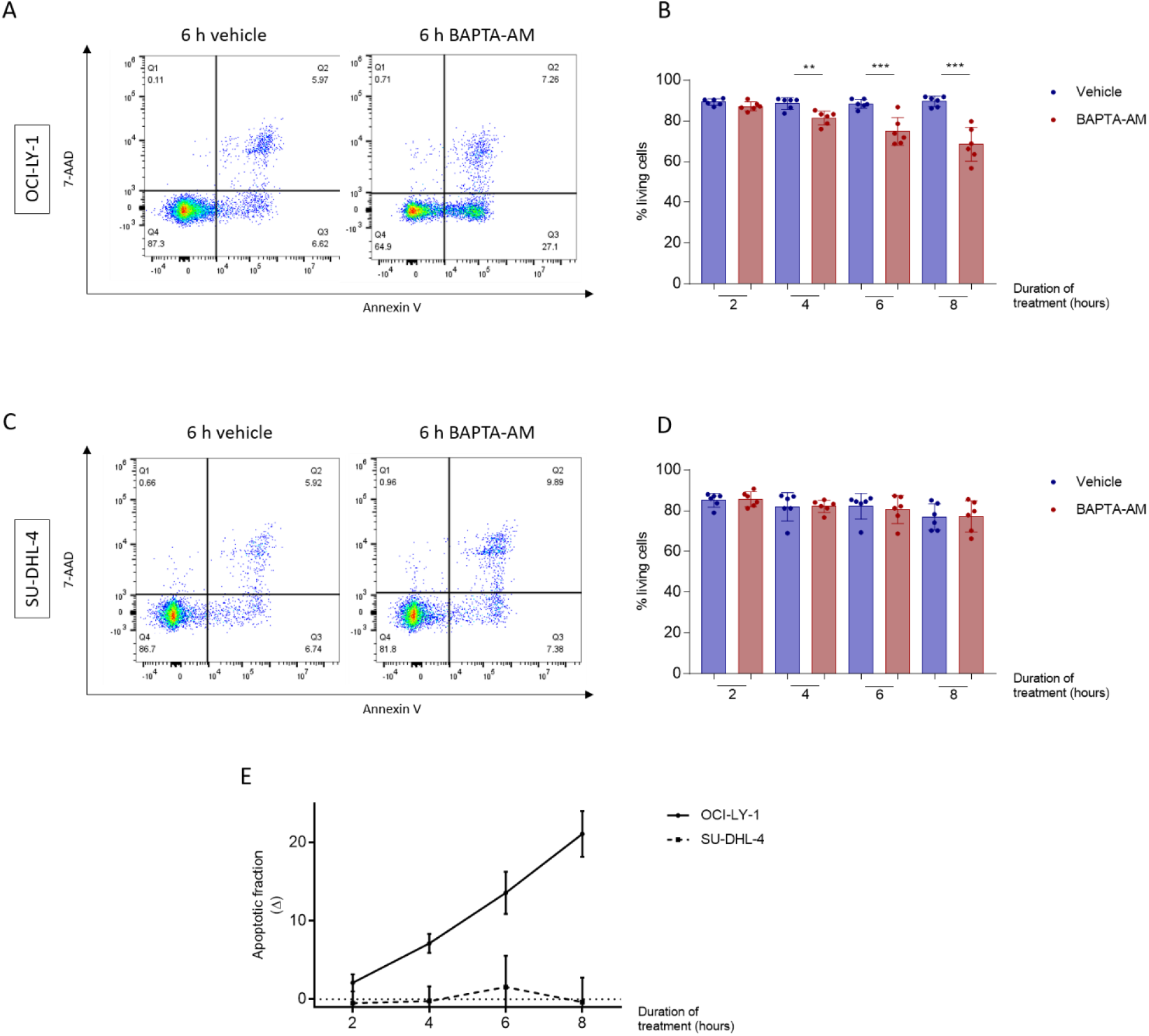
BAPTA-AM-induced apoptosis in OCI-LY-1 and SU-DHL-4 cells. (A-D) Representative graphs and quantitative analysis of apoptosis of OCI-LY-1 (A and B) and SU-DHL-4 (C and D) cells at different time points after the addition of vehicle (DMSO, blue) or 10 μM BAPTA-AM (red). Cells were stained with annexin V-FITC and 7-AAD, and the apoptotic fraction was identified as annexin V-positive cells. Data are represented as the average ± S.D. (N = 5). Statistical significance of differences was determined with a paired ANOVA test. Differences were considered significant when p < 0.05. (** p < 0.01; *** p < 0.001). (E) ΔApoptotic fractions (i.e. % living vehicle-treated cells - % living BAPTA_i_-treated cells) at different time points in OCI-LY-1 and SU-DHL-4 cells evoked by 10 μM BAPTA-AM compared to vehicle. Data are represented as the average ± S.E.M. (N = 6).

### 2. BAPTA_i_ elicits decreased abundance of Mcl-1 in both SU-DHL-4 and OCI-LY-1 cells

DLBCL cells are highly dependent on the expression of anti-apoptotic Bcl-2 family members to counteract apoptosis. We therefore asked whether BAPTA_i_ could differentially impact the expression profile of these proteins in OCI-LY-1 versus SU-DHL-4 cells. Immunoblotting of different Bcl-2 family members in OCI-LY-1 cells and SU-DHL-4 cells treated with BAPTA_i_ for various time periods showed no significant differences in Bcl-2, Bcl-XL and Bim protein levels between vehicle- and BAPTA_i_-treated OCI-LY-1 and SU-DHL-4 cells (Supplementary Figure 1). However, Mcl-1 protein levels rapidly decreased upon BAPTA_i_ treatment in both OCI-LY-1 and SU-DHL-4 cells, with prominent decreases from 4 hours onwards (Figure 3A and B).

**Figure 3:**
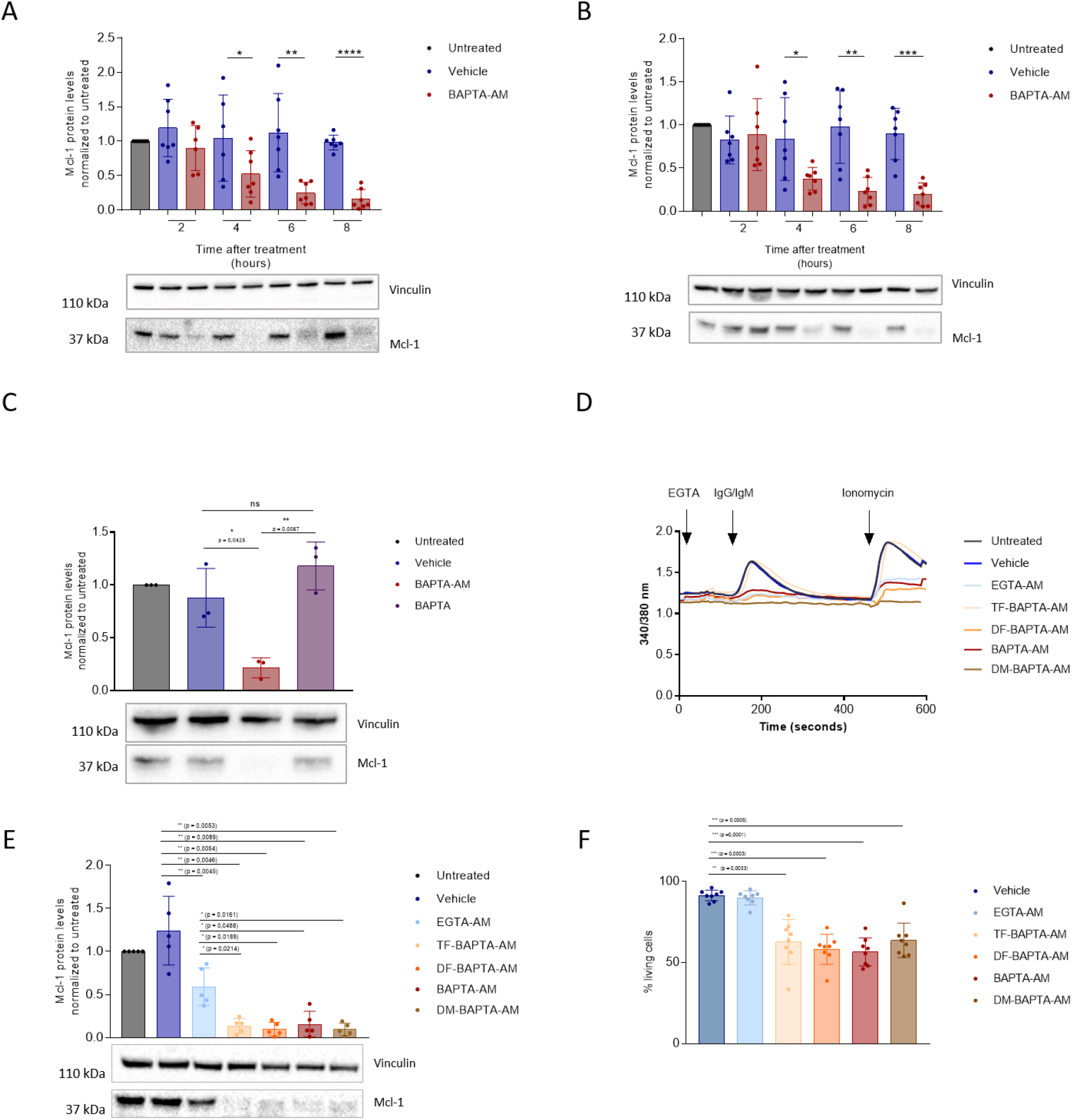
BAPTA-AM caused Mcl-1 downregulation and cell death in a Ca^2+^-independent manner. (A and B) Representative western blot and statistical analysis of Mcl-1 levels normalized to untreated in response to 10 μM of vehicle (dark blue) and BAPTA-AM (red) after 2, 4, 6, 8, and 24 hours in OCI-LY-1 (A) and SU-DHL-4 (B) cells. Vinculin was included as a loading control. Data are represented as the average ± S.D. (N = 7). Statistical significant differences were determined with a paired two-tailed Student’s t-test. Differences were considered significant when p < 0.05. (** p < 0.01; *** p < 0.001). (C) Representative western blot and statistical analysis of Mcl-1 levels normalized to untreated in response to a 6 h treatment with of vehicle (dark blue), 10 μM BAPTA-AM (red) and extracellular BAPTA (purple). Data are represented as the average ± S.D. (N = 3). (D) Mean cytosolic Ca^2+^ measurements using Fura-2-AM in OCI-LY-1 cells after 30 minutes of pretreatment with 10 μM of vehicle (dark blue, N = 8), EGTA-AM (light blue, N = 4), TF-BAPTA-AM (yellow, N = 4), DF-BAPTA-AM (orange, N = 4), BAPTA-AM (red, N = 4) and DM-BAPTA-AM (brown, N = 4). Extracellular EGTA was used to buffer extracellular Ca^2+^, IgG/IgM and ionomycin trigger Ca^2+^ release from the ER and the extracellular space, respectively. (E) Representative western blot and statistical analysis of Mcl-1 levels in response to a 6 h treatment with of vehicle (dark blue), 10 μM EGTA-AM (light blue), TF-BAPTA-AM (yellow), DF-BAPTA-AM (orange), BAPTA-AM (red) and DM-BAPTA-AM (brown). Vinculin was included as a loading control. Data are represented as the average ± S.D. (N = 6). Statistical significant differences were determined with a paired ANOVA test. Differences were considered significant when p < 0.05. (** p < 0.01; *** p < 0.001). (F) Quantitative analysis of apoptosis in OCI-LY-1 in response to 6 h treatment with 10 μM of vehicle, EGTA-AM, TF-BAPTA-AM, DF-BAPTA-AM, BAPTA-AM and DM-BAPTA-AM. Cells were stained with annexin V-FITC and 7-AAD and the apoptotic fraction was identified as annexin V positive cells. Data are represented as the average ± S.D. (N = 8). Statistically significant differences were determined with a paired ANOVA test. Differences were considered significant when p < 0.05. (** p < 0.01; *** p < 0.001).

To ensure that decreased Mcl-1 protein levels were caused by intracellular BAPTA and not extracellular BAPTA, we applied BAPTA as a free acid (not conjugated to acetoxymethyl ester and abbreviated BAPTA_e_ for extracellular BAPTA), which cannot enter the cell. In contrast to BAPTA_i_, BAPTA_e_ failed to provoke a decrease in Mcl-1 protein levels, indicating that the latter effect is indeed caused by intracellularly loaded BAPTA (Figure 3C).

### 3. BAPTA_i_ decreases Mcl-1 protein levels and provokes cell death in a Ca^2+^-independent manner

The above results imply that Ca^2+^ signals, buffered by BAPTA_i_, are essential to maintain adequate Mcl-1 protein levels and thus to sustain the survival of OCI-LY-1 cells that may be dependent on Mcl-1. We therefore determined whether the ability of different BAPTA analogs to chelate intracellular Ca^2+^correlated with their efficacy in decreasing Mcl-1 levels and limit OCI-LY-1 cell survival. Hence, we compared the cell death-inducing properties of BAPTA_i_ in the OCI-LY-1 cell model to three BAPTA_i_ analogs that possess different Ca^2+^-binding affinities (Figure 1). EGTA (loaded into cells as EGTA-AM) was included as a control since it has a Ca^2+^-binding affinity similar to that of BAPTA, albeit with a slower kinetic profile and a different molecular structure, lacking the two benzene rings (Figure 1). Moreover, inclusion of EGTA-AM treatment as a control eliminates any effect caused by the acetoxymethyl (AM) residue after hydrolysis.

First, we validated the distinct Ca^2+^-buffering properties of the different BAPTA_i_ derivatives. We performed cytosolic Ca^2+^ measurements in Fura-2-loaded OCI-LY-1 cells that were exposed to Ca^2+^-mobilizing agonists in the presence of extracellular EGTA to chelate extracellular Ca^2+^. Anti-IgG/IgM activates the B-cell receptor, thereby generating IP_3_ and evoking IP_3_R-mediated Ca^2+^ release from the ER, whereas ionomycin acts as a Ca^2+^ ionophore, thereby mobilizing Ca^2+^ from all internal stores. The various BAPTA analogs affected anti-IgG/IgM- and ionomycin-evoked Ca^2+^ signals in a manner that was consistent with their theoretical K_D_ values for Ca^2+^; DM-BAPTA_i_ was the most potent in buffering the cytosolic Ca^2+^ increases, while TF-BAPTA_i_ did not affect cytosolic Ca^2+^ increases triggered by anti-IgG/IgM and ionomycin (Figure 3D). Hence, in subsequent experiments, TF-BAPTA_i_ was used as a control to determine the role of Ca^2+^ buffering in the cellular effects of BAPTA-AM treatment. After establishing that the BAPTA analogs differentially buffered intracellular Ca^2+^ signals according to their Ca^2+^ affinity, their effects on Mcl-1 levels and cell death in OCI-LY-1 cells were analyzed via western blotting and flow cytometry, respectively (Figure 3E and F). Regardless of their differences in Ca^2+^-buffering capacity, all BAPTA_i_ analogs similarly decreased Mcl-1 protein levels and evoked cell death in OCI-LY-1 cells. Compared to BAPTA_i_ and its different derivatives, EGTAi appeared less effective in reducing Mcl-1 abundance and failed to provoke cell death, although EGTAi buffered the IgG/IgM- and ionomycin-induced Ca^2+^ responses more avidly than TF-BAPTA_i_ and DF-BAPTA_i_. This observation also excludes that the effects of BAPTA_i_ on Mcl-1 and cell death were due to the released AM moiety, since EGTAi, which too releases this AM moiety, does not exert these effects. Finally, we validated that BAPTA_i_ and its derivatives provoked cell death through apoptosis: *i*. Cell death induced by BAPTA_i_ and its derivatives could be rescued by ZVAD-OMe-FMK, a pan-caspase inhibitor (supplemental Figure 2A), and *ii*. BAPTA_i_ and its derivatives provoked the cleavage of PARP, a downstream target of caspase-3. Additionally, in these assays, EGTA_i_ did not have major effects (Supplemental Figure 2B).

### 4. BAPTA_i_-induced cell death is dependent on decreased abundance of Mcl-1

We found that BAPTA_i_ decreased Mcl-1 protein levels both in OCI-LY-1 and SU-DHL-4 cells but mainly provoked cell death in OCI-LY-1 cells. We therefore asked whether the difference in BAPTA_i_ sensitivity could be explained by differences in Mcl-1 dependence between OCI-LY-1 and SU-DHL-4 cells. We exposed both cell lines to varying concentrations of S63845, a high-affinity Mcl-1 antagonist validated to inhibit Mcl-1(12). Furthermore, we included H929 cells, a multiple myeloma cell line known to be highly sensitive to Mcl-1 antagonism. As shown in Figure 4A, S6385 caused a concentration-dependent death of H929 and OCI-LY-1 cells but did not affect SU-DHL-4 cells.

**Figure 4:**
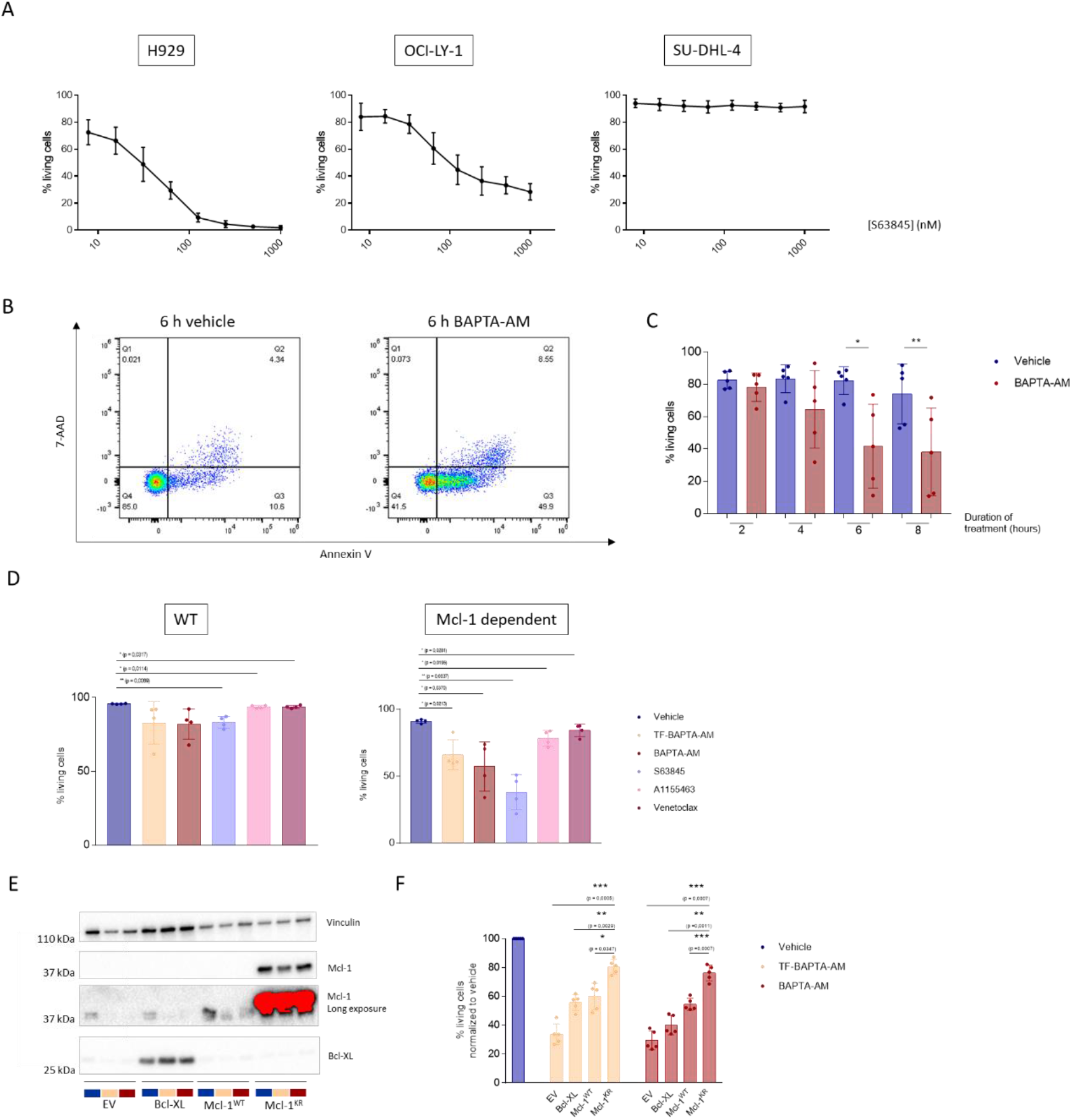
BAPTA_i_-induced cell death is dependent on decreased abundance of Mcl-1 and is prevented by reinforcing Mcl-1 expression. (A) Dose-response curve of the Mcl-1 inhibitor S63845 on cell survival in H929, OCI-LY-1 and SU-DHL-4 cells (left to right) after 6 h of treatment. Cell death was measured using annexin V-FITC and 7-AAD staining, and the apoptotic fraction was identified as annexin V-positive cells. (N = 3). (B & C) Representative graphs and quantitative analysis of apoptosis in H929 cells at different time points after the addition of vehicle (DMSO, blue) or 10 μM BAPTA-AM (red). Cells were stained with annexin V-FITC and 7-AAD, and the apoptotic fraction was identified as annexin V-positive cells. Data are represented as the average ± S.D. (N = 5). Statistical significance of differences was determined with a paired ANOVA test. Differences were considered significant when p < 0.05. (** p < 0.01; *** p < 0.001). (D) SVEC cell lines were treated for 6 hours with vehicle (dark blue), 10 μM TF-BAPTA-AM (yellow), BAPTA-AM (red), Mcl-1 inhibitor S63845 (purple), 1 μM Bcl-XL inhibitor A1155483 (light pink) or 10 μM Bcl-2 inhibitor venetoclax (dark pink). Cells were stained with annexin V-APC and 7-AAD, and cell death was measured via flow cytometry. Data are represented as the average ± S.D. (N = 4). Statistically significant differences were determined with a paired ANOVA test. Differences were considered statistically significant when p < 0.05. (** p < 0.01; *** p < 0.001, **** p < 0.0001). (E) OCI-LY-1 cells were transfected with an empty vector (EV), a Bcl-XL-overexpressing plasmid, a wild-type (WT) Mcl-1-overexpressing plasmid or a plasmid overexpressing a nondegradable KR mutant of Mcl-1. Representative western blot of Mcl-1 and Bcl-XL levels 24 hours after transfection of OCI-LY-1 cells treated for 6 hours with vehicle (dark blue), 10μM TF-BAPTA-AM (yellow) or BAPTA-AM (red) (N = 5). (F) Statistical analysis of cell death in transfected OCI-LY-1 cells after 6 hours of 10 μM TF-BAPTA-AM or BAPTA-AM treatment normalized to vehicle. Cells were stained with annexin V-APC and 7-AAD, and apoptosis was measured via flow cytometry. Data are represented as the average ± S.D. (N = 5). Statistically significant differences were determined with a paired ANOVA test. Differences were considered significant when p < 0.05. (** p < 0.01; *** p < 0.001, **** p < 0.0001).

To underpin the involvement of Mcl-1 in cell death caused by BAPTA_i_ and BAPTA_i_ analogs, we engaged two independent approaches. First, we hypothesized that H929 cells would be sensitive to BAPTA_i_, since they strongly rely on Mcl-1 expression. Consistent with our previous findings, we observed that H929 cells were highly sensitive to BAPTA-AM treatment, and that BAPTA_i_ rapidly provoked cell death in a large subset of H929 cells (Figure 4B and 4C). Second, to ensure that our findings were not due to heterogeneous cellular genetic backgrounds among the different cell lines, we implemented an isogenic cell system using cells with genetically engineered dependence towards Mcl-1(23). ‘Mito-priming’ is based on the equimolar co-expression of pro- and anti-apoptotic Bcl-2 proteins to artificially generate an addiction to a specific anti-apoptotic Bcl-2 protein. We compared the effects of BAPTA_i_ and BAPTA_i_ analogs on the survival of wild-type (WT, parental control) and Mcl-1-dependent SVECs (Figure 4D). Each cell line was treated with TF-BAPTA-AM and BAPTA-AM, while their Mcl-1 dependence was benchmarked using specific antagonists for Mcl-1 (S63845), Bcl-2 (venetoclax) and Bcl-XL (A1155483). SVECs engineered to be dependent on Mcl-1 died in response to S63845 but not to venetoclax or A1155483 treatment, while the parental SVEC control controls were much less affected by all three BH3 mimetics. Both BAPTA_i_ and TF-BAPTA_i_ provoked significant cell death in Mcl-1-dependent SVEC cells but were only marginally effective in the WT parental control cells. These results indicate that BAPTA_i_-induced cell death depends on cell addiction to Mcl-1 for survival and is not a cell type-or DLBCL-specific phenomenon.

The data presented above indicate that treating cells with BAPTA-AM or BAPTA-AM analogs leads to a rapid loss of Mcl-1 proteins within cells. Since Mcl-1 is a short-lived and rapidly turned-over protein, we examined whether enhancing Mcl-1 protein levels could alleviate sensitivity to BAPTA_i_. To this end, we transfected OCI-LY-1 cells with a nondegradable Mcl-1 mutant (Mcl-1^K/R^)(24). This Mcl-1 mutant was obtained through sequential mutagenesis of the established ubiquitination sites in human Mcl-1 (amino acids 5, 40, 136, 194, and 197) from lysine to arginine to prevent its ubiquitination and subsequent proteasomal degradation. To allow appropriate estimation of the effectiveness of the nondegradable Mcl-1^K/R^ mutant, parallel cultures were transfected with either an empty vector or a vector overexpressing wild-type Mcl-1. We also overexpressed Bcl-XL to further confirm the specific mediation of the effects by Mcl-1 rather than just any Bcl-2-family member. Overexpression of Bcl-XL, wild-type Mcl-1 and the nondegradable Mcl-1^K/R^ mutant in OCI-LY-1 cells was confirmed by immunoblotting (Figure 4E, lanes 4, 7 and 10 versus lane 1). Immunoblot analysis showed that wild-type Mcl-1 levels were strongly reduced in BAPTA-AM-treated cells. This was the case for both endogenous (empty vector, EV) and ectopically expressed Mcl-1 (Mcl-1^WT^). In contrast, both Bcl-XL and the nondegradable Mcl-1^K/R^ mutant protein levels were virtually unaffected by treatment of OCI-LY-1 cells with BAPTA-AM or TF-BAPTA-AM. Ectopic expression of the nondegradable Mcl-1^K/R^ mutant conferred significant protection against apoptosis towards treatment with BAPTA-AM or TF-BAPTA-AM compared to cells transfected with an empty vector, Bcl-XL or WT Mcl-1. These results further indicate that BAPTA_i_ triggered cell death in OCI-LY-1 cells by decreasing Mcl-1 protein levels, since restoring Mcl-1 protein levels could counteract this process.

### 5. BAPTA_i_ reduces Mcl-1 protein levels by suppressing mTORC1, an important driver of *MCL-1* translation

Next, we examined the mechanism by which BAPTA_i_ evoked a decline in Mcl-1 protein levels. We first analyzed the effect of BAPTA_i_ and its analogs on *MCL-1* mRNA levels but did not observe any change following treatment with the different BAPTA-AM compounds, indicating the absence of effects on *MCL-1* transcription (Figure 5A). To determine whether BAPTA_i_ enhanced Mcl-1 protein degradation, we pretreated OCI-LY-1 cells with cycloheximide to inhibit the *de novo* translation of Mcl-1 (Figure 5B). Since Mcl-1 is rapidly turned over by proteasomal degradation, halting translation using cycloheximide results in a rapid decline in Mcl-1 (Figure 5B, blue bars). However, compared to the vehicle control, neither BAPTA_i_ nor TF-BAPTA_i_ accelerated the cycloheximide-induced decline in Mcl-1 protein levels (Figure 5B, yellow and red bars). Thus, BAPTA_i_ and TF-BAPTA_i_ did not promote Mcl-1 degradation.

**Figure 5:**
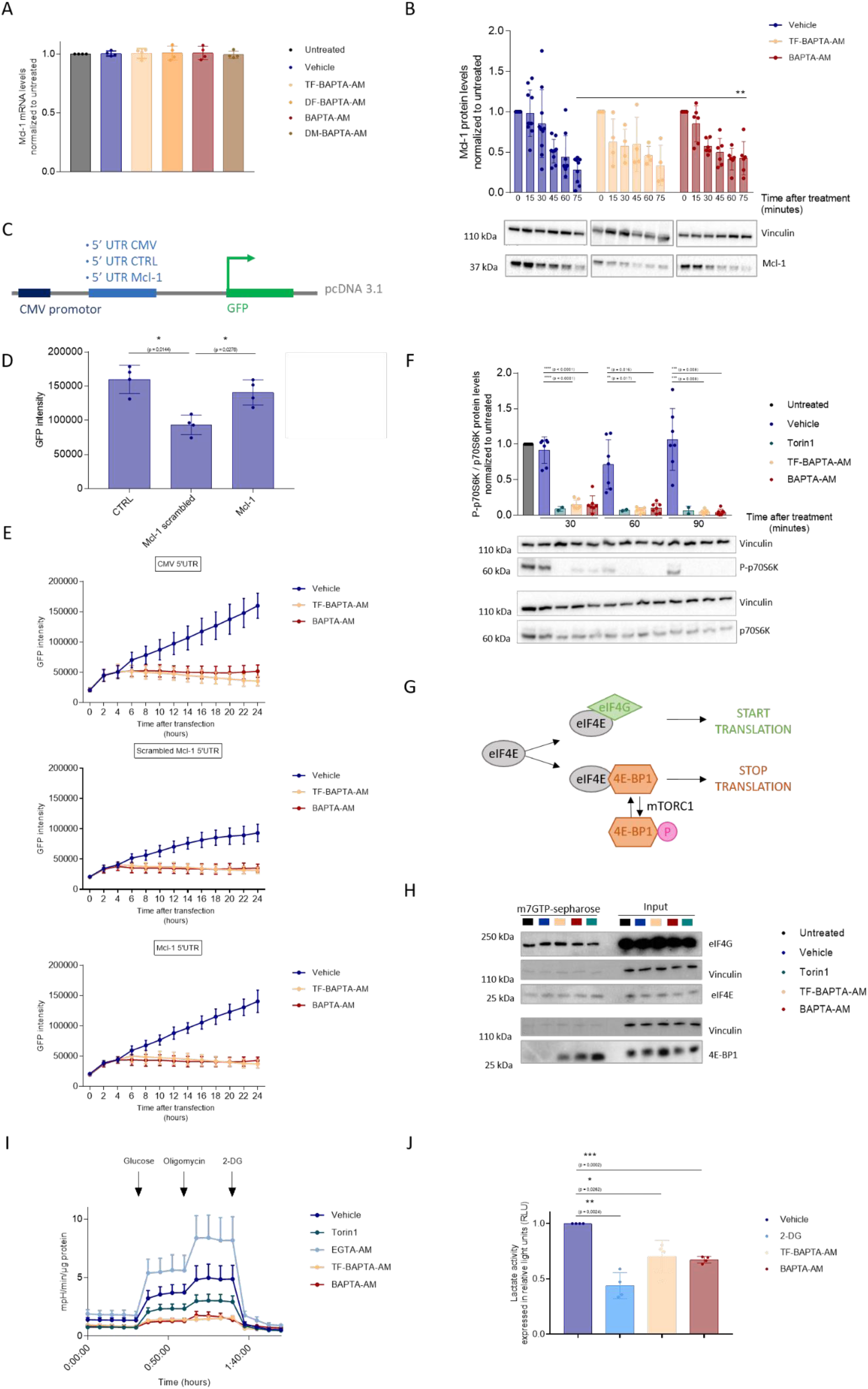
BAPTA_i_ inhibits translational activity by hampering glycolysis. (A) Statistical analysis of Mcl-1 mRNA levels after 6 h treatment with 10 μM vehicle (dark blue), TF-BAPTA-AM (yellow), DF-BAPTA-AM (orange), BAPTA-AM (red) and DM-BAPTA-AM (brown) normalized to untreated (black) Data are represented as the average ± S.D. (N = 4). Statistically significant differences were determined with a paired ANOVA test. Differences were considered significant when p < 0.05. (** p < 0.01; *** p < 0.001). (B) Representative western blot and densitometric analysis of Mcl-1 protein levels after 0, 15, 30, 45, 60 and 75 minutes of vehicle (dark blue, N = 10), 10 μM TF-BAPTA-AM (yellow, N = 4) and BAPTA-AM (red, N = 6) treatment. Prior to this, the cells were pretreated with 20 μg/mL cycloheximide. Quantified Mcl-1 levels were normalized to the loading control (vinculin) and calculated relative to the untreated condition. (C) Schematic representation of the constructs used in 4D and E. A pcDNA3.1 reporter vector was inserted with the original (Mcl-1) or a scrambled (control, CTRL) 5’UTR region of Mcl-1. A construct with the original CMV 5’ UTR was included as an additional control. OCI-LY-1 cells were subsequently electroporated with one of the constructs, and GFP intensity was measured. (D) GFP intensity levels after 24 hours of transfection with the aforementioned constructs and vehicle treatment. Data are represented as the average ± S.D. (N = 4) Statistically significant differences were determined with paired ANOVA. Differences were considered significant when p < 0.05. (** p < 0.01; *** p < 0.001). (E) Graphical representation of GFP intensity after transfection with one of the aforementioned constructs. After 4 hours, transfected cells were treated with 10 μM of the caspase inhibitor ZVAD-OMe-FMK and 30 minutes later with 10 μM vehicle (dark blue), TF-BAPTA-AM (yellow) or BAPTA-AM (red). Measurement of GFP intensity was continued for 24 hours (N = 4). (F) Representative western blot and densitometric analysis of P-p706K/total p70S6K normalized to untreated cells after 30, 60 and 90 minutes of treatment with vehicle (dark blue, N = 7), 2 μM Torin1 (green, N = 2), 10 μM TF-BAPTA-AM (yellow, N = 7) or BAPTA-AM (red, N = 7). Quantified (P-)p70S6K levels were normalized to the loading control (vinculin) and calculated relative to the untreated condition. Data are represented as the average ± S.D. Statistically significant differences were determined with a paired ANOVA test. Differences were considered significant when p < 0.05. (** p < 0.01; *** p < 0.001). (G) Translation initiation factor eIF4E forms active complexes with eIF4G (green), leading to mRNA recruitment. eIF4G competes with 4E-BP1 (orange), which forms inactive complexes with eIF4E, leading to inhibition of translation. Phosphorylation of 4E-BP1 by mTORC1 causes its release from eIF4E. (H) Western blot of protein extracts from OCI-LY-1 cells treated for 90 minutes with vehicle (dark blue), 10 μM TF-BAPTA-AM (yellow), BAPTA-AM (red) and 2 μM Torin1 (green). Lysates were incubated with m7GTP-sepharose beads, and bound proteins were denatured and subjected to western blotting to reveal eIF4E bound to eIF4G (active complexes) or to 4E-BP1 (inactive complexes). Vinculin was included as a loading control (N = 3). (I) OCI-LY-1 cells were pretreated for 1 h with vehicle (dark blue, N = 8), 1 μM Torin1 (green, N = 5), 10 μM EGTA-AM (light blue, N = 3), TF-BAPTA-AM (yellow, N = 8) or BAPTA-AM (red, N = 8) in glucose-free medium. All experimental extracellular pH values were measured after the addition of glucose (normal glycolytic activity), oligomycin (maximal glycolytic activity) and 2-DG (nonglycolytic acidification). In each experiment, data were obtained in three-fold and normalized to the amount of protein. Data are represented as the average ± S.E.M. (J) OCI-LY-1 cells were pretreated for 1 h with 10 μM of vehicle (dark blue), 10 mM 2-DG (bright blue), 10 μM TF-BAPTA-AM (yellow) and 10 μM BAPTA-AM (red). Extracellular lactate was measured and the amount of extracellular lactate correlates with absorption levels expressed in relative light units (RLU) normalized to vehicle treatment. Data are presented as the average ± S.E.M. (N = 4).

Since BAPTA_i_ neither inhibited *MCL-1* transcription nor accelerated Mcl-1 degradation, we assessed whether BAPTA_i_ could suppress Mcl-1 translation. We monitored translation by cloning the 5’ untranslated region (UTR) of *MCL-1* in front of an open reading frame encoding GFP (Figure 5C). We also cloned the 5’ UTR of CMV to monitor general translation. These constructs were benchmarked against a control that lacked a 5’ UTR (for this, we used a scrambled version of the 5’ UTR of *MCL-1*), which provides the background translation in our system, potentially due to leaky translational events. First, we validated that the 5’ UTR of *MCL-1* and CMV significantly augmented the translation of GFP compared to the scrambled 5’ UTR of *MCL-1* (Figure 5D). Then, we assessed the impact of BAPTA_i_ and TF-BAPTA_i_ on both *MCL-1* and CMV 5’ UTRs to determine their effect on Mcl-1 translation and general translation events. *MCL-1* 5’ UTR- and CMV 5’ UTR-driven translation was decreased by BAPTA_i_ and TF-BAPTA_i_, indicating that BAPTA_i_ inhibits Mcl-1 translation and CMV-driven translation in a Ca^2+^-independent manner (Figure 5E).

Since Mcl-1 translation is driven by mammalian target of rapamycin complex 1 (mTORC1), we assessed the effect of BAPTA_i_ on the phosphorylation of p70S6K, a downstream target of mTORC1 signaling that drives mRNA translation and controls cell growth and metabolism (Figure 5F)(25). OCI-LY-1 cells were treated with vehicle, TF-BAPTA-AM or BAPTA-AM. The selective and high-affinity mTORC1 inhibitor Torin1 was included as a positive control. Samples were taken after 30, 60 and 90 minutes, and the ratio of P-p70S6K/total p70S6K was analyzed via western blot. All treatments similarly and significantly lowered the phosphorylation of p70S6K, indicating that BAPTA_i_ suppresses mTORC1 activity in a Ca^2+^-independent manner. Moreover, these results indicate that BAPTA_i_ may diminish general protein translation by inhibiting mTORC1 activity.

To further strengthen the conclusion that BAPTA_i_ inhibits mTORC1-driven protein translation, we examined eIF4E, the rate-limiting regulator of cap-dependent mRNA translation. eIF4E either forms a complex with eIF4G, thereby recruiting mRNA and driving translation, or forms an inactive complex with 4E-BP1, thereby competing with eIF4G for binding eIF4E and limiting translation. Phosphorylation of 4E-BP1 by mTORC1 causes its release from eIF4E to allow cap-dependent translation to proceed (Figure 5G). Thus, the eIF4E-binding ratio of eIF4G/4E-BP1 provides a molecular readout for mTORC1-driven mRNA translation activity. Therefore, we pulled-down eIF4E using m7GTP-sepharose beads and checked the binding of eIF4G and 4E-BP1 in lysates of OCI-LY-1 cells treated for 90 minutes with TF-BAPTA-AM, BAPTA-AM or Torin1 as a control (Figure 5H). All treatments elicited a prominent increase in 4E-BP1 binding to eIF4E compared to the vehicle control, indicating that BAPTA_i_ indeed inhibits cap-dependent mRNA translation and that this occurs independently of its Ca^2+^-chelating properties.

### 6. BAPTA_i_ rapidly suppresses glycolysis in cells preceding cell death by abrogating the conversion of fructose-6-phosphate into fructose-1,6-bisphosphate

Since mTORC1 activity is tightly controlled by the metabolic state of the cell, we monitored glycolytic activity in OCI-LY-1 and SU-DHL-4 cells exposed to TF-BAPTA_i_ and BAPTA_i_. To eliminate any effect caused by the AM residue or by mTOR inhibition itself, EGTA_i_ and Torin1 were included as controls. Using a Seahorse extracellular flux analyzer, we analyzed the change in extracellular pH (extracellular acidification rate, ECAR), which is an approximate measure of glycolytic activity. Although BAPTA_i_-induced cell death was only observed after approximately 4 hours with OCI-LY-1 cells (Figure 2B), pretreatment with the pan-caspase inhibitor ZVAD-OMe-FMK was performed to avoid any potential adverse effects of cell death initiation. Within 1 hour, BAPTA_i_ and TF-BAPTA_i_ strongly lowered ECAR, while Torin1, EGTA_i_ or vehicle did not exert such an effect (Figure 5I). In addition, we performed extracellular lactate measurements to validate that both BAPTA_i_ and TF-BAPTA_i_ lowered extracellular lactate levels (Figure 5J).

Next, to unravel the mechanism by which BAPTA_i_ interferes with glycolysis at the molecular level, we set up a metabolomics approach using nonradioactive ^13^C glucose (Figure 6). We cultured OCI-LY-1 cells in the presence of U-^13^C-labeled glucose, so that the cells could incorporate the stable ^13^C isotopes into their metabolic pathways. With mass spectrometry, the abundance and labeling patterns (isotopologs and fractional contribution) of metabolites could be detected, allowing us to assess the activity and connectivity of metabolic pathways. The cells were treated for one hour with vehicle, BAPTA-AM or TF-BAPTA-AM. BAPTA_i_ caused a significant decrease in fructose 1,6-bisphosphate levels, which also occurred in response to TF-BAPTA_i_ (Figure 6). Downstream metabolites, but not upstream metabolites, were also reduced. This suggests that BAPTA_i_ provokes Ca^2+^-independent inhibition of glycolysis at the level of fructose 6-phosphate conversion into fructose 1,6-bisphosphate. Glycolytic metabolites downstream of fructose 1,6-bisphosphate were less reduced by TF-BAPTA_i_ than BAPTA_i_ (Figure 6). This could be caused by better maintained activity of the pentose phosphate pathway (PPP), which converts fructose 6-phosphate into glyceraldehyde 3-phosphate that can re-enter the glycolytic pathway, by-passing the block.

**Figure 6:**
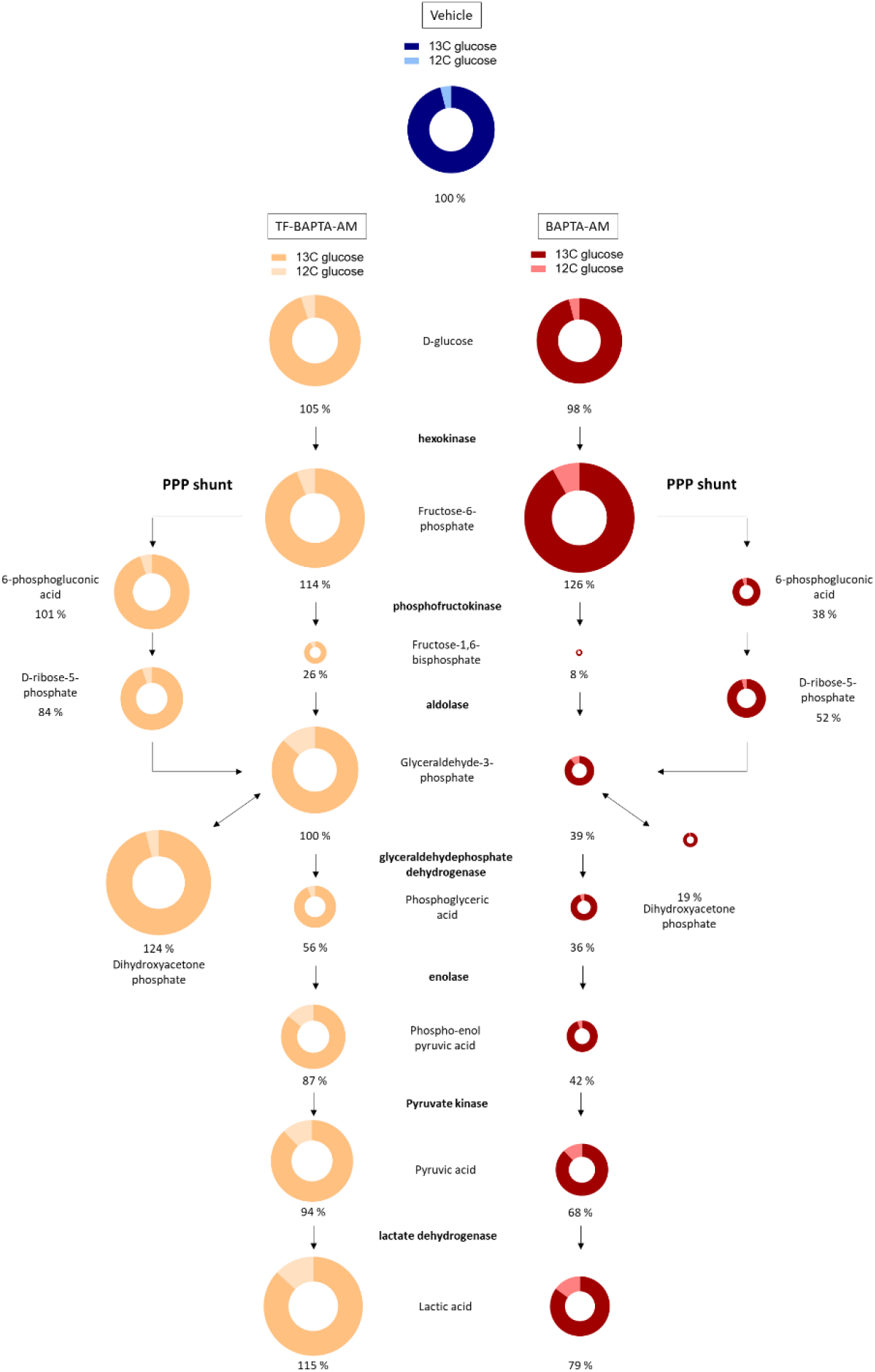
BAPTA_i_ and TF-BAPTA_i_ potently inhibit the conversion of fructose 1-phosphate into fructose 1,6-bisphosphate. Twenty-four hours prior to treatment, OCI-LY-1 cells were cultured in glucose-free IMDM supplemented with ^13^C glucose. Afterwards, the cells were subsequently treated for 1 h with vehicle (dark blue), 10 μM TF-BAPTA-AM (yellow) or BAPTA-AM (red). Abundances and fractional contributions of ^12^C vs ^13^C for each glycolytic metabolite were measured using mass spectrometry. The size of the doughnut represents the abundance of each metabolite normalized to the vehicle condition; % changes in abundance are indicated beneath each doughnut. The color of the doughnut represents the relative contributions of ^13^C (dark) and ^12^C (light) (N = 3).

### 7. BAPTA directly inhibits the activity of purified, recombinantly expressed PFKFB3

The production of fructose 1,6-bisphosphate is catalyzed by the phosphofructokinase 1 (PFK1) enzyme, whose activity strongly depends on fructose 2,6-bisphosphate, a metabolite produced by the activity of the 6-phosphofructo-2-kinase/fructose-2,6-bisphosphatase 3 (PFKFB3) enzyme. We therefore asked whether BAPTA could directly impact the master regulator PFKFB3. We measured the kinase activity of the recombinant human PFKFB3 enzyme using a cell-free biochemical assay based on measuring the production of ADP in the presence of different concentrations of BAPTA (ranging from 1 μM to 1 mM). This concentration range was chosen since previous studies indicated that application of 10 μM BAPTA-AM in the extracellular environment (also used in experiments in our work) resulted in the accumulation of BAPTA in the cytosol up to 1-2 mM (15). In this assay, BAPTA directly inhibited PFKFB3 activity with an IC_50_ of 110 μM (Figure 7A). Appropriate controls (counter assay, see Material and Methods) confirmed that BAPTA did not affect the ATP-depleting and luciferase detection enzymes included in the ADP-Glo system used in the kinase assay (not shown). EGTA, which has a Ca^2+^-binding affinity similar to that of BAPTA but a different molecular structure, did not affect PFKFB3 kinase activity (Figure 7B). Next, we asked whether the inhibition of PFKFB3 by BAPTA was influenced by Ca^2+^. We found that increasing the Ca^2+^ concentrations in the assay buffer alleviated the inhibitory effect of BAPTA (applied at 1 mM) on PFKFB3 (while this had no impact on PFKFB3 activity in the presence of EGTA), further indicating that the inhibition of PFKFB3 by BAPTA occurs through its Ca^2+^-free form (Figure 7C). It’s interesting to note that adding 0.5 mM Ca^2+^ to 1 mM BAPTA thus anticipating ~50%Ca^2+^-free BAPTA and ~50% Ca^2+^-bound BAPTA corresponds to about a 50% reduction of the inhibitory effect of BAPTA. Saturating BAPTA with Ca^2+^ by adding 1 or 2 mM Ca^2+^ to 1 mM BAPTA completely alleviated the inhibitory effect of BAPTA on PFKFB3. This shows that BAPTA can directly inhibit PFKFB3 activity in its Ca^2+^-free form, but not in its Ca^2+^-complexed form.

**Figure 7:**
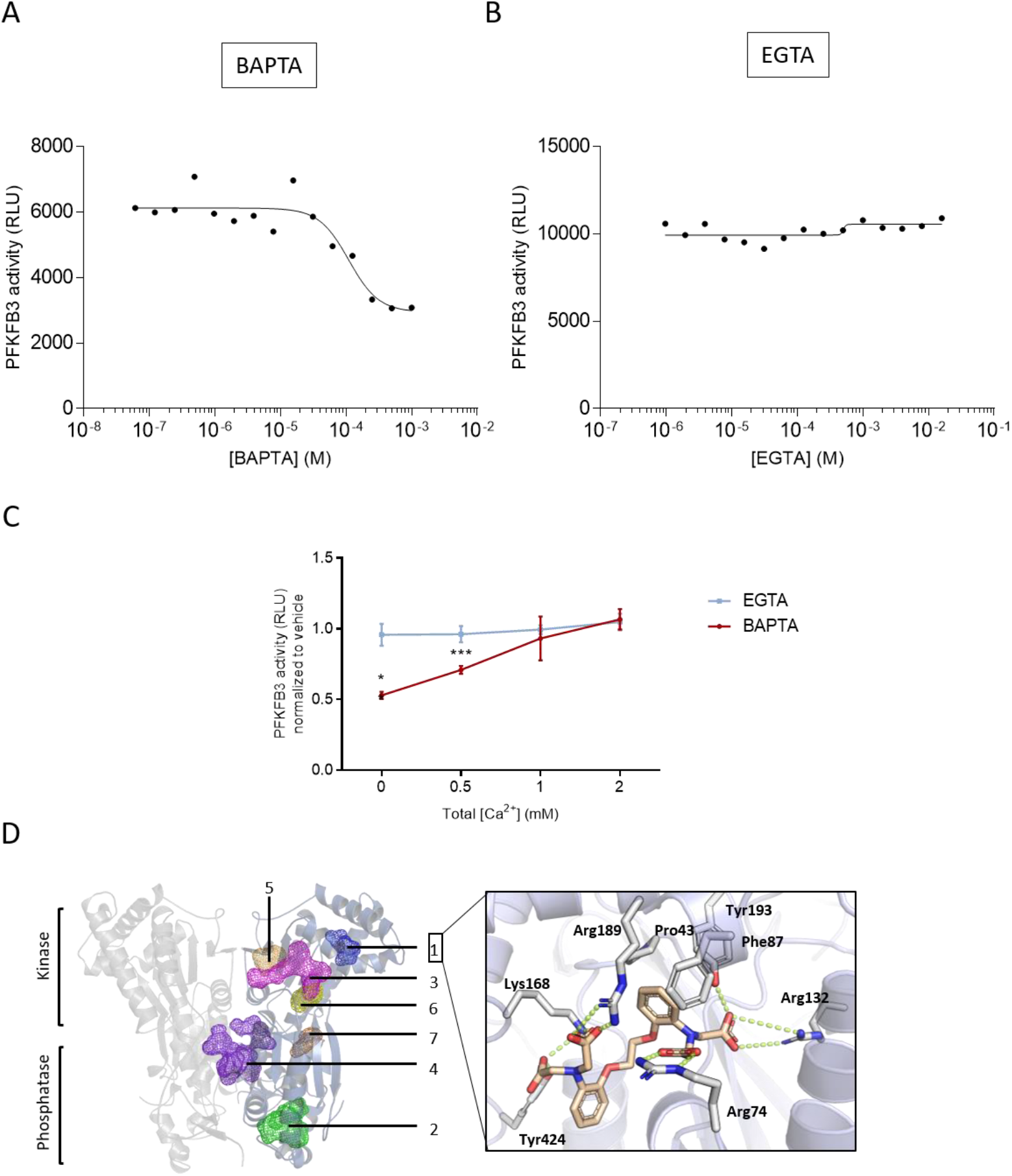
BAPTA directly interacts with PFKFB3, leading to inhibition of its kinase activity. (A & B) In vitro measurement of PFKFB3 activity in the presence of increasing concentrations of EGTA (A) and BAPTA (B). (C) In vitro measurement of PFKFB3 activity in the presence of 1 mM BAPTA (red) or 1 mM EGTA (blue) normalized to vehicle (H2O) conditions, measured with increasing Ca^2+^concentrations (0 to 2 mM CaCl2) in the assay buffer. (D) The PFKFB3 dimer is represented in two different shades of cartoon. FTmap was used to identify 7 putative pockets, which have been mapped on one monomer, including the fructose-6-phosphate site, (1), the fructose-2,6 bisphosphate site (2), the ATP binding site (3), a dimerization site (4) and other minor cavities (5–7). BAPTA was docked to each of the binding pockets leading to 7 Goldscores: 83.9, 94.8, 67.3, 74.1, 64.8, 58.4, and 63.7 from pockets 1 to 7, respectively. The highest scoring pockets were further investigated using induced fit docking yielding a putative BAPTA binding mode to pocket 1 of the X site. BAPTA makes four salt bridges between its four carboxylate functional groups, and Arg 74, Arg 132, Lys 168, and Arg189 are further stabilized by hydrogen bonds with Tyr193 and Tyr424. The aromatic ring is buried between the hydrophobic side chains of Pro43, Phe87 and Tyr193.

To further substantiate that BAPTA can directly impact PFKFB3, we performed a molecular modeling approach by docking BAPTA into the different putative binding sites (Figure 7D). We identified seven different sites, including the ATP-binding site and the fructose 6-phosphate sites in the kinase domain and the fructose 2,6-bisphosphate site in the hydrolase domain and a site at the dimerization interface. The docking simulations preferred a binding mode to both fructose 6-phosphate binding sites bound in the kinase and hydrolase domains (with scores of 94.8 and 83.9 over scoring 74.1 to 58.4 in the other domains). This is not surprising, as both domains are hallmarked by a series of positively charged residues to coordinate multiple phosphates. To obtain a more detailed view, induced-fit docking was performed at both sites, hypothesizing that the acid functionalities of BAPTA mimic the fructose phosphate groups. This resulted in a preferred binding mode of BAPTA to the fructose 6-phosphate sites within the kinase domain (−10.8 kcal/mol) over the hydrolase site (−9.3 kcal/mol). With EGTA, this dropped to −9.0 kcal/mol. The docking simulations suggest specific binding of BAPTA to the fructose 6-phosphate substrate-binding site in the kinase domain competing with the substrate and ATP binding site via phosphate-mimicking interactions.

### 8. PFKFB3 inhibition suppresses mTORC1 activity, thereby evoking a decline in Mcl-1 protein levels

Finally, we sought to establish a causal link between the inhibition of PFKFB3 activity, inhibition of mTORC1 activity and a subsequent decrease of Mcl-1 protein levels in OCI-LY-1 and SU-DHL-4 cells, thereby mimicking the effect of BAPTA_i_. We therefore used the AZ PFKFB3 67 compound, a PFKFB3 inhibitor(26). We exposed OCI-LY-1 and SU-DHL-4 cells to 25 μM AZ PFKFB3 67, a concentration limiting PFKFB3 activity in cells(27). Similar to Torin1 and BAPTA_i_, AZ PFKFB3 67 decreased the phospho-p70S6K/total p70S6K ratio and reduced Mcl-1 levels (Figure 8). These results indicate that pharmacological inhibition of PFKFB3 is sufficient to suppress mTORC1 activity and downstream signaling events, thereby resulting in a decline in Mcl-1 protein levels.

**Figure 8:**
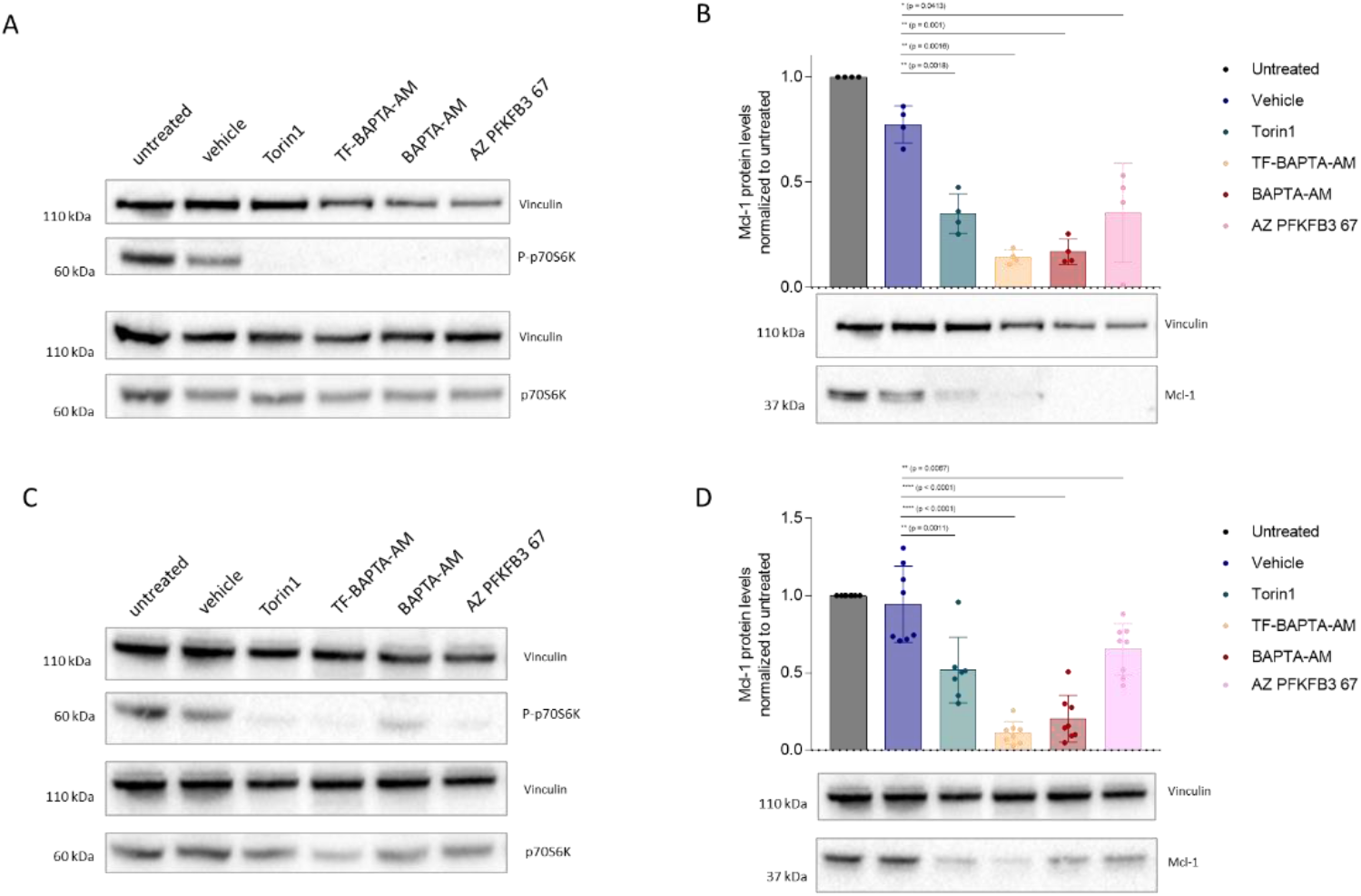
Pharmacological inhibition of PFKFB3 by BAPTA leads to dephosphorylation of p70S6K and decreased abundance of Mcl-1. (A - D) SU-DHL-4 (A & B) and OCI-LY-1 (C & D) cells were treated with vehicle, 2 μM Torin1, 10 μM TF-BAPTA-AM, 10 μM BAPTA-AM or 25 μM PFKFB3 inhibitor (AZ PFKFB3 67). (A & C) Representative western blots of P-p70s6K and p70S6K. (N = 3). (B & D) Representative western blot and densitometric analysis of Mcl-1 levels in response to 6 h of treatment with vehicle (dark blue), 2 μM Torin1 (dark green), 10 μM TF-BAPTA-AM (yellow), 10 μM BAPTA-AM (red) or 25 μM PFKFB3 inhibitor (pink). Quantified Mcl-1 levels were normalized to the loading control (vinculin) and calculated relative to the untreated condition. Data are represented as the average ± S.D. (N = 4 (B), N = 8 (D)). Statistically significant differences were determined with a paired ANOVA test. Differences were considered significant when p < 0.05. (** p < 0.01; *** p < 0.001).

## Discussion

Our results reveal an unprecedented and Ca^2+^-independent effect of BAPTA_i_, a high-affinity, rapid Ca^2+^-chelating agent (see Figure 9 for a simplified model). We demonstrate that BAPTA_i_ hampers the conversion of fructose 6-phosphate into fructose 1,6-bisphosphate by directly inhibiting the activity of PFKFB3, a major regulatory enzyme of this step. The inhibition of PFKFB3 by BAPTA occurred via its Ca^2+^-free form. Consequently, BAPTA_i_ strongly suppressed mTORC1 activity, thereby abrogating downstream mTORC1-controlled events such as protein translation. Although likely overall mTORC1-driven protein translation is suppressed, BAPTA_i_ particularly affects the protein levels of short-lived proteins such as anti-apoptotic Mcl-1, whose protein levels rapidly decline upon inhibition of mTORC1. As a consequence, in cell models in which Mcl-1 is critical for cell survival, BAPTA_i_ evokes cell death. Thus, our work revealed novel intracellular crosstalk where PFKFB3 inhibition reduces Mcl-1 protein levels by impeding mTORC1 activity.

**Figure 9:**
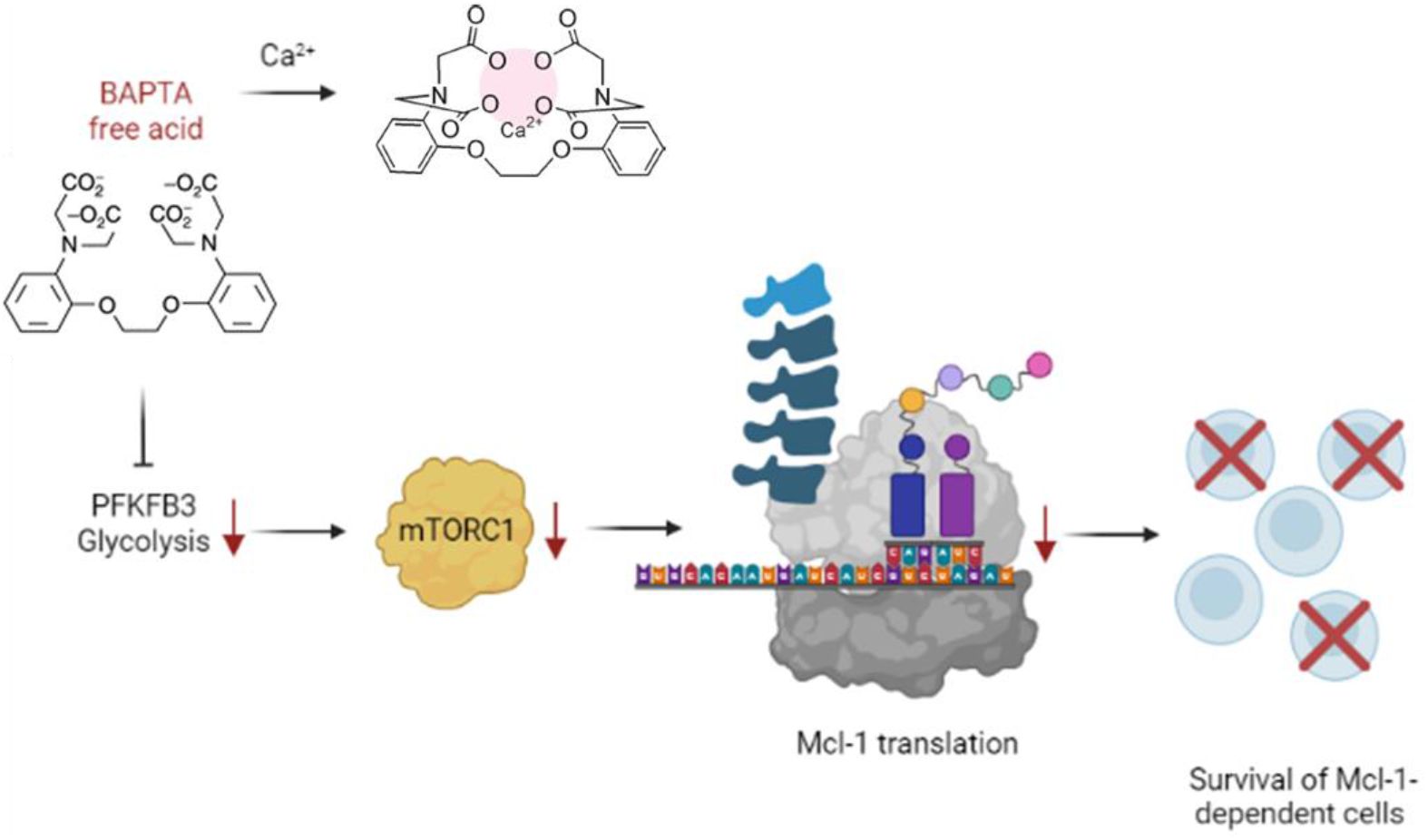
Proposed model. BAPTA, independent of its Ca^2+^-chelating properties and in its Ca^2+^-free form, directly inhibits PFKFB3 activity. This suppresses mTORC1 activity, an important driver of protein translation, including anti-apoptotic Mcl-1 proteins, which are short-lived proteins rapidly turned over by the proteasome. As a consequence, loss of mTORC1 activity evokes a rapid decline in Mcl-1 protein levels. In cell models in which Mcl-1 is critical for survival such as Mcl-1-dependent DLBCL cells, this will result in cell death.

We previously demonstrated that treatment with BAPTA_i_ could boost the cell death response of DLBCL cells to venetoclax, an FDA-approved Bcl-2 antagonist (14). This is particularly interesting since resistance to venetoclax treatment is emerging(14),(28),(29). The ability of BAPTA_i_ to enhance the cell death response of cancer cells to BH3 mimetics has also been observed in other studies. For instance, ovarian carcinoma cells were sensitized to Bcl-XL inhibition upon BAPTA-AM treatment (20). As BAPTA_i_ is a high-affinity buffer of cytosolic Ca^2+^, these findings suggested a yet unappreciated interplay between constitutive Ca^2+^ signaling, anticipated to be suppressed by BAPTA_i_, and the anti-apoptotic function of the Bcl-2 family. However, no studies so far have performed controls using BAPTA_i_ variants with low Ca^2+^ affinity such as TF-BAPTA_i_ that cannot affect basal cytosolic Ca^2+^ levels. Here, we demonstrate that BAPTA_i_ can exert profound cellular effects that are completely independent of its Ca^2+^-chelating properties.

Using an unbiased ^13^C tracer metabolomics approach, we found that BAPTA_i_ impedes a proximal step in glycolysis, namely, the conversion of fructose 6-phosphate into fructose 1,6-bisphosphate. We postulate that BAPTA_i_ evokes a direct inhibition of PFKFB3, as BAPTA can inhibit the activity of purified PFKFB3 enzyme in *in vitro* activity assays. In these assays, the inhibition of PFKFB3 by BAPTA was characterized by an IC_50_ of 110 μM. Although this might seem a high concentration, it is important to note that previous work indicated that extracellular application of 10 μM BAPTA-AM results in cytosolic BAPTA concentrations of about 1 to 2 mM (15), thus well-above the presumed IC_50_, enabling *in cellulo* inhibition of PFKFB3 by BAPTA_i_. As a consequence of this PFKFB3 inhibition, we found that downstream pathways such as mTORC1 signaling, a major driver of protein translation, were impaired. This statement is supported by pharmacological PFKFB3 inhibitors that could mimic BAPTA_i_ and inhibit mTORC1 activity. Following mTORC1 inhibition, protein translation is suppressed including the translation of anti-apoptotic Mcl-1 proteins, thereby rapidly decreasing its protein levels. Mcl-1 protein levels are exquisitely susceptible to impaired translation since Mcl-1 is a short-lived protein that is quickly turned over through proteasomal degradation. When cells are addicted to Mcl-1 for their survival, they become sensitive to decreases in Mcl-1 protein levels and thus to BAPTA-AM treatment. This implies that BAPTA_i_ does not elicit a general, off-target toxicity but provokes a specific Ca^2+^-independent inhibition of glycolysis.

Mcl-1 is an important pharmacological target since it is an emergent resistance factor in tumor cells treated with BH3 mimetics, such as ABT-199 (venetoclax), ABT-263 (navitoclax) and ABT-737. The need for therapeutic agents targeting Mcl-1 is therefore acute(11). Accordingly, specific Mcl-1 inhibitors have been developed, showing promising results as single agents as well as in combination with other BH3 mimetics(28)–(32). However, increased abundance of nontargeted Bcl-2 family members is not the only resistance mechanism in tumor cells. Following prolonged treatment with venetoclax, several studies reported mutations in the hydrophobic groove of Bcl-2, decreasing the affinity of venetoclax for Bcl-2 and limiting its therapeutic effect(33),(34). Although few studies have addressed this concern so far, it is plausible that a similar mechanism would occur for Mcl-1 when treating tumor cells with a BH3-mimetic Mcl-1 inhibitor. It could therefore be beneficial to target Mcl-1 at the level of its transcription, translation, or degradation. Indeed, it has been shown that inhibition of the mTORC1 complex, and by extension Mcl-1 translation, shows antitumor effects in lymphoma as a single agent and in combination with venetoclax(35),(36). This is supported by our own results demonstrating that BAPTA-induced inhibition of mTORC1 and consequent reduction of Mcl-1 protein levels leads to cell death in Mcl-1-dependent lymphoma cell lines. mTORC1 is an energy sensing complex that controls anabolic processes, including ribosome biogenesis and synthesis of nucleotides, proteins and lipids, and inhibits catabolism (autophagy)(25). While mTORC1 aligns energy supplies with anabolic and catabolic activities in physiological conditions, in cancer cells it lies at the core of metabolic reprogramming to increase proliferation. Accordingly, mTORC1 inhibitors, so-called rapalogs, have been extensively evaluated as potential therapeutics in cancer treatments(37).

Our findings are in line with other studies demonstrating a link between glycolysis, mTORC1 activity and mRNA translation. Indeed, the team of Adams and coworkers revealed that 2-deoxyglucose (2-DG), which inhibits the first step in the glycolytic pathway, caused a decrease in p70S6K phosphorylation and an increase in 4E-BP1 activity, indicating that inhibition of glycolysis using 2-DG provoked a decrease in general translation activity(38). In the same study, 2-DG rapidly affected steady-state Mcl-1 levels, sensitizing human hematopoietic tumor cell lines to ABT-737, a nonselective Bcl-2/Bcl-XL-antagonizing BH3 mimetic.

Although regulation of mTORC1 by amino acids is well established, the exact mechanisms by which glucose controls mTORC1 activity remain poorly understood. Different mechanisms may contribute to glucose sensing by mTORC1, including the AMPK, ULK1/LARS and PFKFB3 pathways(39). PFKFB3 is a kinase enzyme that catalyzes the phosphorylation of fructose 6-phosphate into fructose 2,6-bisphosphate and acts as a stimulator of PFK1, one of the rate-limiting enzymes of glycolysis. PFKFB3 controls the translocation of mTORC1 to lysosomes by direct interaction with Rag B and subsequent mTORC1 activity(40). Our data reveal that PFKFB3 activity sustains mTORC1 activity and thus mTORC1 signaling. These findings open up future research avenues since increased abundances of PFKFB3 are found in several cancers, and has been associated with cancer hallmarks such as carcinogenesis, cancer cell proliferation, drug resistance and the tumor microenvironment.(41),(42).

Inhibition of glycolytic activity then decreases mTORC1 activity, leading to impaired (Mcl-1) translation and cancer cell death. Indeed, cancer cells are often characterized by a high rate of glycolytic flux, known as the Warburg effect. This mechanism allows fast ATP production and generates carbon sources for *de novo* nucleotide synthesis; both mechanisms enabling rapid proliferation(43),(44). Direct inhibition of glycolysis has therefore been extensively explored as an anticancer strategy(45), underpinning the importance of the elucidated noncanonical effects of BAPTA_i_.

BAPTA has been applied in more than a thousand publications as a“proof” that Ca^2+^ signaling is essential for certain biological processes. For effects of BAPTA_i_ to be fully ascribed to chelation of Ca^2+^, experimental protocols should incorporate the use of adequate controls such as a TF-BAPTA_i_. Our work also expands the previous research performed by Smith and colleagues. Their data revealed that several Ca^2+^-chelating agents and fluorescent Ca^2+^ indicators, including BAPTA, could elicit Ca^2+^-independent effects, such as inhibition of Na^+^/K^+^-ATPase, an ion pump critical for maintenance of ionic balances and control cell volume(15),(16). In the future, it will be interesting to assess a proteome-wide view of the distinct targets of intracellular BAPTA. Such studies are needed to discern effects related to BAPTA’s Ca^2+^-chelating properties versus its potential direct binding to targets and biochemically altering their activity.

In conclusion, BAPTA elicits important cellular effects independent of its ability to chelate Ca^2+^. We found that BAPTA inhibited the kinase activity of the glycolytic stimulator PFKFB3, leading to impaired glycolysis. This resulted in impairment of mTORC1 activity leading to dysfunctional translation. Consequently, the cellular expression of Mcl-1, a short-lived anti-apoptotic protein quickly affected by the lack of *de novo* translation, was reduced, promoting cell death in Mcl-1-dependent cancer cells.

## Materials and methods

### 1. Cell culture and transfections

SU-DHL-4 and OCI-LY-1 DLBCL cell lines were kindly provided by Dr. Anthony Letai (Dana Farber Cancer Institute, Boston, Massachusetts, USA). H929 cells were purchased from DSMZ (Braunschweig, Germany). OCI-LY-1 cells were cultured in suspension in Iscove modified Dulbecco’s medium (IMDM, Life Technologies, Brussels, Belgium). SU-DHL-4 and H929 cells were cultured in suspension in Roswell Park Memorial Institute (RPMI-1640) medium (Life Technologies, Brussels, Belgium). Mito-primed seminal vesicle epithelial cells (SVECs) were kindly obtained from Prof. Stephen Tait.(23) All media were supplemented with 10% fetal bovine serum (FBS, Life Technologies), 2% GlutaMAX™ Supplement (Life Technologies, Brussels, Belgium) and 2% penicillin/streptomycin (Life Technologies, Brussels, Belgium). Cultures were incubated at 37°C and 5% CO_2_, and sterile conditions were maintained at all times. Cells were validated through STR profiling and were cultured in mycoplasma-free conditions, whereby cell cultures were monitored once every two weeks for mycoplasma infection. Research with human cell lines was approved by ethical committee UZ Leuven (S63808).

### 2. Transfection

Twenty-four hours after seeding, OCI-LY-1 cells were transfected using the Amaxa^®^ Cell Line Nucleofector^®^ Kit L (Lonza, Basel, Switzerland), program C-05 as previously described(46). Cells were briefly transfected with the constructs described below and collected at 18 hours posttransfection to use in experiments and to confirm transfection via western blot.

### 3. Reagents, antibodies and constructs

The following reagents were used in this study. EGTA (Acros Organics, Geel, Belgium, 409910250), dimethyl sulfoxide (DMSO, Sigma-Aldrich, Overijse, Belgium), Fura-2-AM (Life Technologies, Carlsbad, CA, USA, F1221), 2-DG (Sigma-Aldrich, Overijse, Belgium; purity ≥ 98 %), 1,2-Bis-(o-Aminophenoxy)-ethane-N,N,N’,N’-tetraacetic acid, tetraacetoxymethyl ester (BAPTA-AM, Life Technologies, Brussels, Belgium), 5,5’,6,6’-Tetrafluoro-BAPTA-AM (Interchim, Montluçon, France), 5,5’-difluoro-BAPTA-AM and 5,5’-dimethyl-BAPTA-AM (Sigma Aldrich, Overijse, Belgium), S63845 (Gentaur, Kampenhout, Belgium), venetoclax (ABT-199, Active Biochem, Kowloon, Hong Kong), A1155463 (Selleck Chemicals, Houston, USA), Z-Val-Ala-DL-Asp(OMe)-fluoromethylketone (ZVAD-(OMe)-FMK, ABCAM, Cambridge, UK) cycloheximide (C7698, Sigma-Aldrich, Overijse, Belgium), AZ PFKFB3 67 (Bio-techne, Abingdon, UK).

In addition to BAPTA itself, three different BAPTA analogs were used in this study: tetrafluoro-BAPTA (TF-BAPTA), difluoro-BAPTA (DF-BAPTA) and dimethyl-BAPTA (DM-BAPTA) (Figure 1). The value of EGTA reported is obtained at pH 7.4 and 20 °C. Whereas placing fluor groups on the benzene ring of BAPTA (para-position, or meta- and para-positions, for DF-BAPTA and TF-BAPTA, respectively) severely reduces the affinity for binding Ca^2+^, methyl groups (para position) augment the Ca^2+^-chelating properties of BAPTA. All of the Ca^2+^ chelators used in this study were introduced into the cells as acetoxymethyl esters using standard loading procedures.

The following primary antibodies were used in this study: anti-Mcl-1 (4572, Cell Signaling Technology), anti-Bcl-XL (MA5-15142, Invitrogen), anti-Bcl-2 HRP (sc-7382, Santa Cruz), anti-vinculin (V9131, Sigma Aldrich), anti-P-p70S6K (9234S, Cell Signaling Technology), anti-p70S6K (9202S, Cell Signaling Technology), anti-GFP, anti-PARP (9542S, Cell Signaling Technology), and anti-β actin (A5441, Sigma Aldrich).

The pcDNA3.1 vector bearing the sequence of the nondegradable human gene Mcl-1 with mutated ubiquitination sites, referred to as Mcl-1^K/R^ was kindly provided by Professor Marc Diederich(47). Corresponding empty control plasmids were used in parallel. pcDNA3.1-hMcl-1 was a gift from Roger Davis (Addgene plasmid 25375, Cambridge, MA, USA) and is indicated in the text as WT Mcl-1. Bcl-XL overexpression was achieved using a pcDNA3.1(+) vector encoding human Bcl-XL.

The Mcl-1 5’UTR sequence inserted into the pcDNA3.1 plasmid to generate a GFP reporter construct was GCGGCCGCGCAACCCTCCGGAAGCTGCCGCCCCTTTCCCCTTTTATCGGAATACTTTTTTTAAAAAAAAAGAGTTCGCTGGCGCCACCCCGTAGGACTGGCCGCCCTAAAAGTGATAAAGGAGCTGCTCGCCACTTCTCACTTCCGCTTCCTTCCAGTAAGGAGTCGGGGTCTTCCCCAGTTTTCTCAGCCAGGCGGCGGACTGGCAGAATTC.A scrambled Mcl-1 5’UTR sequence served as a negative control: GCGGCCGCAGTTTTTAGTACAGCAGCCCCCCATAACGGCGCCGCCTAGCGCTCAGTAGTCTTTTAGGCGTTGGGAACAGTGCTGCACGATAGGGTCGTCTCCAGCGGGCCATTGTTGCATACACCATACCGCTGGCGTTATCCCTGTGCATCCGGGCTCATCCGCCAACCTGGTACCACTAGCATCTTATCCCAAAGGGCCGACCATTTCCCACGGAATTC.

### 4. Apoptosis assay

Cells (5 × 10^5^ cells/ml) were treated as indicated in the Results, pelleted by centrifugation, and incubated with annexin V-FITC/7-AAD or annexin V-APC in the presence of annexin V binding buffer. Cell suspensions were analyzed with an Attune^®^ Acoustic Focusing Flow Cytometer (Applied Biosystems). Cell death by apoptosis was scored by quantifying the population of annexin V-FITC-positive cells (blue laser; BL-1) or annexin V-APC-positive cells (red laser; RL-1). Flow cytometric data were plotted and analyzed using FlowJo software (version 10).

### 5. RNA extraction and PCR analysis

Cells were harvested and centrifuged for 5 minutes at 500 × g. RNA was extracted using the HighPure RNA Isolation kit (Roche, Mannheim, Germany; # 11828665001) according to the manufacturer’s protocol. cDNA was prepared using the High Capacity cDNA Reverse Transcription kit (Applied Biosystems, Brussels, Belgium; # 4368814) according to the manufacturer’s protocol. mRNA was amplified using GoTaq Green master mix (Promega, Leiden, The Netherlands; # M7112) and specific primers for the mRNA of interest (IDT, Leuven, Belgium) and separated on a 2.5% Ultrapure agarose (Invitrogen, # 16500-500) gel containing 0.005% EtBr (Invitrogen, # 15585-011).

### 6. Western blot analysis

Cells were washed with phosphate-buffered saline and incubated at 4°C with lysis buffer (20 mM Tris-HCl (pH 7.5), 150 mM NaCl, 1.5 mM MgCl2, 0.5 mM dithiothreitol, 1% Triton-X-100, and one tablet of complete EDTA-free protease inhibitor (Thermo Scientific, Brussels, Belgium)) for 30 minutes. Cell lysates were centrifuged for 5 minutes at 12000 × g and analyzed by Western blotting as previously described(48). Western blot quantification was performed using Image Lab 5.2 software.

### 7. Cytosolic Ca^2+^ measurements

OCI-LY-1 cells were seeded in poly-L-lysine-coated 96-well plates (Greiner Bio One, Vilvoorde, Belgium) at a density of 5 × 10^5^ cells/ml. The cells were loaded for 30 minutes with 1.25 μM Fura-2 AM at 25 °C in modified Krebs solution, followed by a 30-minute treatment with the compounds of interest. Fluorescence was monitored on a FlexStation 3 microplate reader (Molecular Devices, Sunnyvale, CA, USA) by alternately exciting the Ca^2+^ indicator at 340 and 380 nm and collecting emitted fluorescence above 510 nm, as described previously(49).

### 8. Live cell imaging

OCI-LY-1 cells (5 × 10^6^) were transfected with a pcDNA3.1 plasmid bearing either the CMV 5’ UTR, a scrambled Mcl-1 5’ UTR or the original 5’ UTR. All vectors expressed GFP, and both cell confluence and GFP intensity were measured. Following transfection, the cells were incubated in the IncuCyte^®^ Live Cell Analyzer, and microscopic pictures (Nikon 10x objective) were taken every two hours. After four hours, the cells were treated with 10 μM pan-caspase inhibitor ZVAD-OMe-FMK and 30 minutes later with 10 μM vehicle, TF-BAPTA-AM or BAPTA-AM. The cell plate was subsequently placed in IncuCyte^®^ for another 20 hours.

### 9. Metabolic flux analysis

Glycolysis was measured with the Seahorse Glycolysis Stress Test on a Seahorse XFe24 Analyzer (Agilent Technologies, Heverlee, Belgium), which determines the extracellular acidification rate (ECAR) as a measure of glycolytic activity. OCI-LY-1 cells (5 × 10^5^ cells/ml) were pretreated for 1 hour in Seahorse Seahorse XF base medium supplemented with glutamine (103334-100, Agilent Technologies, Heverlee, Belgium) in a CO_2_-free incubator. After pretreatment, the ECAR was measured after the subsequent addition of glucose (10 mM final concentration), oligomycin (1 μM final concentration) and 2-deoxyglucose (2-DG, 50 mM final concentration) to assess normal glycolytic activity, maximal glycolytic activity and the nonglycolytic acidification level, respectively. Afterwards, protein concentrations in each well were measured using the BCA assay and used for normalization.

### 10. Extracellular lactate assay

OCI-LY-1 cells (5 × 10^5^ cells/ml) were washed twice in prewarmed PBS and resuspended in Seahorse XF base medium supplemented with glutamine (103334-100, Agilent Technologies, Heverlee, Belgium). Cells were treated with compounds of interest as indicated, and glucose (10 mM) was added 30 minutes after treatment. Extracellular lactate was measured according to the manufacturer’s protocol using the Lactate-Glo^™^ Assay kit (J5021, Promega, Leiden, The Netherlands).

### 11. Tracer metabolomics analysis

OCI-LY-1 cells were seeded at 3 × 10^5^ cells/ml in IMDM without glucose (AL230A, HiMedia, Mumbai, India) and supplemented with 4.5 mg/ml ^13^C or ^12^C glucose, 10% dialyzed FBS (A3382001, Thermo Scientific, Brussels, Belgium) and 2% penicillin/streptomycin (Life Technologies, Brussels, Belgium). After 24 hours, the cells were treated with vehicle or the indicated compound. One hour later, the cells were centrifuged at 1500 x g for 5 minutes at 4°C, washed with 1 ml of ice-cold NaCl (150 mM in H2O) and again centrifuged under the same conditions. Subsequently, the washing solution was removed, and 150 μL of extraction buffer (80% MeOH at −80°C) was added using a precooled pipet tip. Cells were vortexed until the pellet was completely dissolved. The solution was centrifuged at 20 000 x g for 15 minutes at 4 °C, the supernatant was used for further analysis, and the protein pellet was used to measure the protein concentration (BCA assay).

Ten microliters of each supernatant sample was loaded into a Dionex UltiMate 3000 LC System (Thermo Scientific Bremen, Germany) equipped with a C-18 column (Acquity UPLC-HSS T3 1. 8 μm; 2.1 × 150 mm, Waters) coupled to a Q Exactive Orbitrap mass spectrometer (Thermo Scientific) operating in negative ion mode. A step gradient was carried out using solvent A (10 mM TBA and 15 mM acetic acid) and solvent B (100% MeOH). The gradient started with 0% solvent B and 100% solvent A and remained at 0% B until 2 minutes post injection. A linear gradient to 37% B was carried out until 7 minutes and increased to 41% until 14 minutes. Between 14 and 26 minutes, the gradient increased to 100% B and remained at 100% B for 4 minutes. At 30 minutes, the gradient returned to 0% B. The chromatography was stopped at 40 minutes. The flow was kept constant at 250 μL/min, and the column was placed at 25°C throughout the analysis. The MS operated in full scan mode (m/z range: [70 −1050]) using a spray voltage of 3.2 kV, capillary temperature of 320°C, sheath gas at 10.0, and auxiliary gas at 5.0. The AGC target was set at 3e6 using a resolution of 140.000, with a maximum IT fill time of 512 ms. Data collection was performed using Xcalibur software (Thermo Scientific). The data analysis was performed by integrating the peak areas (El-Maven - Polly - Elucidata).

### 12. PFKFB3 kinase activity assay

Biochemical Assay Principle: PFKFB3 enzyme activity was estimated by calculating the amount of ADP generated in a kinase reaction. ADP generation was measured using an ADP Glo kit (Promega, Leiden, The Netherlands) with no deviation from the recommended protocol.

Kinase activity assay: PFKFB3 kinase activity was measured according to a published protocol with slight modifications.(26) Briefly, 2X base buffer was prepared as a standard for all kinase reactions. This buffer contained 100 mM HEPES (pH 7.5), 200 mM KCl, 10 mM MgCl_2_, 8 mM dithiothreitol, 0.02% Triton X100, 0.02% BSA and 4 mM fructose 6-phosphate. Immediately prior to starting the assay, kinase enzyme was added to the base buffer at a 2X concentration of 40 nM. Each well with PFKFB3 enzyme received 2 μL of the 2X enzyme/base buffer solution. Compounds of interest (1 μL) were added, followed by a 30-minute preincubation. Next, 1 μL containing 80 ATP μM was added (giving a final concentration of 20 μM ATP, 2 mM fructose 6-phosphate and 20 mM enzyme) to start the reaction. The kinase reaction was stopped after two hours by adding 4 μL of ADP-Glo Reagent. One hour later, 8 μL of ADP-Glo Detection Reagent was added. After an additional hour, a PerkinElmer Envision^®^ with an enhanced luminescence module was used to measure the luminescence signal generated in each well. For background subtraction, wells receiving ATP but enzyme-free base buffer and without compound addition were used. To verify the potential effect of compounds on the ADP-Glo kit enzymes (counter assay), wells with compounds received enzyme-free base buffer.

### 13. Molecular docking

The PFKFB3 protein structure in dimeric form was retrieved from the PDB (3qpv)(50) and prepared for docking using the protonate3D functionality implemented in MOE (Molecular Operating Environment (MOE), 2020.09 Chemical Computing Group ULC, 1010 Sherbooke St. West, Suite #910, Montreal, QC, Canada, H3A 2R7, 2022.). The BAPTA (and EGTA) ligand was also modeled in MOE using the mmff94x forcefield as a deprotonated ligand. FTmap was used to identify the potential ligand binding sites both in the monomer and at the dimer interface. Each site was next used for docking of the BAPTA ligand using GOLD with standard parameters.(51) The docking score was calculated using the GOLDfitness score. Induced fit docking was performed on the top scoring pockets using MOE, and the free energy of binding of the ligand to the receptor was calculated using the GBVI/WSA method (Molecular Operating Environment (MOE), 2020.09 Chemical Computing Group ULC, 1010 Sherbooke St. West, Suite #910, Montreal, QC, Canada, H3A 2R7, 2022).

### 14. Statistical analysis

All statistical tests were performed using Prism 7 (GraphPad, La Jolla, CA, USA). Two-group comparisons were made using Student’s *t*-test assuming equal variances. Multiple groups were analyzed by one-way ANOVA with Greenhouse-Geisser corrections and *P*-values were included. Unless otherwise indicated, all data are presented as the mean ± S.D. with a significant *P*-value (* *P* < 0.05, ** *P* < 0.01, *** *P* < 0.001, **** *P* < 0.0001).

## Data availability

All relevant data are presented in the manuscript.

## Acknowledgements

The authors wish to thank Anja Florizoone and Rita La Rovere for excellent technical help. The research was funded by grants from the Research Foundation – Flanders (G090118N, G0E7520N, G081821N and G094522N to GB) and from the Research Council – KU Leuven (C14/19/099 and AKUL/19/34 to GB). GB and MDB are part of the FWO Scientific Research Network CaSign (W0.019.17N and W0.014.22N). We thank Dr. Marc Diederich for sharing the Mcl-1^K/R^-expression plasmid.

## Author contributions

Performing experiments: FS, KW, AS, AV, GE, MD, FSR. Analyzing results: FS, BG. Discussion: FS, MK, MD, MDB. Design research: FS, MDB, GB. Supervision and conceptualization: GB. Acquired funding: GB. Provided critical reagents: SWT

## References

1. Berridge, M., Lipp, P. & Bootman, M. The versatility and universality of calcium signalling. Nat. Rev. Mol. Cell Biol. 1, 11–21 (2000).

2. Clapham, D. Calcium signaling. Cell 131, 1047–1058 (2007).

3. Monteith, G., Prevarskaya, N. & Roberts-Thomson, S. The calcium-cancer signalling nexus. Nat. Rev. Cancer 17, 367–380 (2017).

4. Marchi, S. & Pinton, P. Alterations of calcium homeostasis in cancer cells. Curr. Opin. Pharmacol. 29, 1–6 (2016).

5. Akl, H. & Bultynck, G. Altered Ca2+ signaling in cancer cells: Proto-oncogenes and tumor suppressors targeting IP3 receptors. Biochim. Biophys. Acta - Rev. Cancer 1835, 180–193 (2013).

6. Ando, H., Kawaai, K., Bonneau, B. & Mikoshiba, K. Remodeling of Ca 2+ signaling in cancer: Regulation of inositol 1,4,5-trisphosphate receptors through oncogenes and tumor suppressors. Adv. Biol. Regul. 68, 64–76 (2018).

7. Grimm, S. The ER-mitochondria interface: the social network of cell death. Biochim. Biophys. Acta 1823, 327–334 (2012).

8. Rosa, N., Ivanova, H., Wagner, L. E., Kale, J., La Rovere, R., Welkenhuyzen, K., Louros, N., Karamanou, S., Shabardina, V., Lemmens, I., Vandermarliere, E., Hamada, K., Ando, H., Rousseau, F., Schymkowitz, J., Tavernier, J., Mikoshiba, K.,… Bultynck, G. Bcl-xL acts as an inhibitor of IP 3 R channels, thereby antagonizing Ca 2+-driven apoptosis. Cell Death Differ. 29, 788–805 (2022).

9. Sarosiek, K., Ni Chonghaile, T. & Letai, A. Mitochondria: gatekeepers of response to chemotherapy. Trends Cell Biol. 23, 612–619 (2013).

10. Peperzak, V., Slinger, E., Ter Burg, J. & Eldering, E. Functional disparities among BCL-2 members in tonsillar and leukemic B-cell subsets assessed by BH3-mimetic profiling. Cell Death Differ. 24, 111–119 (2017).

11. Bose, P., Gandhi, V. & Konopleva, M. Pathways and mechanisms of venetoclax resistance. Leuk. Lymphoma 58, 2026–2039 (2017).

12. Villalobos-Ortiz, M., Ryan, J., Mashaka, T. N., Opferman, J. T. & Letai, A. BH3 profiling discriminates on-target small molecule BH3 mimetics from putative mimetics. Cell Death Differ. 27, 999 (2020).

13. Kehr, S. & Vogler, M. It’s time to die: BH3 mimetics in solid tumors. Biochim. Biophys. acta. Mol. cell Res. 1868, (2021).

14. Vervloessem, T., Ivanova, H., Luyten, T., Parys, J. B. & Bultynck, G. The selective Bcl-2 inhibitor venetoclax, a BH3 mimetic, does not dysregulate intracellular Ca2+ signaling. Biochim. Biophys. Acta - Mol. Cell Res. 1864, 968–976 (2017).

15. Smith, N. A., Kress, B. T., Lu, Y., Chandler-Militello, D., Benraiss, A. & Nedergaard, M. Fluorescent Ca2+ indicators directly inhibit the Na,K-ATPase and disrupt cellular functions. Sci. Signal. 11, (2018).

16. Bootman, M. D., Allman, S., Rietdorf, K. & Bultynck, G. Deleterious effects of calcium indicators within cells; an inconvenient truth. Cell Calcium vol. 73 82–87 (2018).

17. Turner, C., Connell, J., Blackstone, K. & Ringler, S. Loss of calcium and increased apoptosis within the same neuron. Brain Res. 1128, 50–60 (2007).

18. Williams, J., Hou, Y., Ni, H. & Ding, W. Role of intracellular calcium in proteasome inhibitor-induced endoplasmic reticulum stress, autophagy, and cell death. Pharm. Res. 30, 2279–2289 (2013).

19. Zhang, L., Cheng, X., Xu, S., Bao, J. & Yu, H. Curcumin induces endoplasmic reticulum stress-associated apoptosis in human papillary thyroid carcinoma BCPAP cells via disruption of intracellular calcium homeostasis. Medicine (Baltimore). 97, (2018).

20. Bonnefond, M.-L., Lambert, B., Giffard, F., Abeilard, E., Brotin, E., Louis, M.-H., Gueye, M. S., Gauduchon, P., Poulain, L. & N’Diaye, M. Calcium signals inhibition sensitizes ovarian carcinoma cells to anti-Bcl-xL strategies through Mcl-1 down-regulation. Apoptosis 2015 204 20, 535–550 (2015).

21. Saoudi, Y., Rousseau, B., Doussière, J., Charrasse, S., Gauthier-Rouvière, C., Morin, N., Sautet-Laugier, C., Denarier, E., Scaïfe, R., Mioskowski, C. & Job, D. Calcium-independent cytoskeleton disassembly induced by BAPTA. Eur. J. Biochem. 271, 3255–3264 (2004).

22. Furuta, A., Tanaka, M., Omata, W., Nagasawa, M., Kojima, I. & Shibata, H. Microtubule disruption with BAPTA and dimethyl BAPTA by a calcium chelation-independent mechanism in 3T3-L1 adipocytes. Endocr. J. 56, 235–243 (2009).

23. Lopez, J., Bessou, M., Riley, J. S., Giampazolias, E., Todt, F., Rochegüe, T., Oberst, A., Green, D. R., Edlich, F., Ichim, G. & Tait, S. W. G. Mito-priming as a method to engineer Bcl-2 addiction. Nat. Commun. 7, (2016).

24. Masuoka, H., Mott, J., Bronk, S., Werneburg, N., Akazawa, Y., Kaufmann, S. & Gores, G. Mcl-1 degradation during hepatocyte lipoapoptosis. J. Biol. Chem. 284, 30039–30048 (2009).

25. XM, M. & J, B. Molecular mechanisms of mTOR-mediated translational control. Nat. Rev. Mol. Cell Biol. 10, 307–318 (2009).

26. Boyd, S., Brookfield, J. L., Critchlow, S. E., Cumming, I. A., Curtis, N. J., Debreczeni, J., Degorce, S. L., Donald, C., Evans, N. J., Groombridge, S., Hopcroft, P., Jones, N. P., Kettle, J. G., Lamont, S., Lewis, H. J., MacFaull, P., McLoughlin, S. B.,… Wingfield, J. Structure-Based Design of Potent and Selective Inhibitors of the Metabolic Kinase PFKFB3. J. Med. Chem. 58, 3611–3625 (2015).

27. Savva, C., Gonz Alez-Granillo, M., Li, X., Angelin, B., Korach-Andr, M., Modder, M., Kuipers, E. N., Held, N. M., In, W., Panhuis, H., Ruppert, P. M. M., Kersten, S., Kooijman, S., Guisas, B., Houtkooper, R., Rensen, P., Boon, M.,… De Meyer, G. 3PO inhibits glycolysis but does not bind to 6-phosphofructo-2-kinase/fructose-2,6-bisphosphatase-3 (PFKFB3). Atherosclerosis 315, e87 (2020).

28. Ramsey, H. E., Fischer, M. A., Lee, T., Gorska, A. E., Arrate, M. P., Fuller, L., Boyd, K. L., Strickland, S. A., Sensintaffar, J., Hogdal, L. J., Ayers, G. D., Olejniczak, E. T., Fesik, S. W. & Savona, M. R. A Novel MCL-1 Inhibitor Combined with Venetoclax Rescues Venetoclax-Resistant Acute Myelogenous Leukemia. Cancer Discov. 8, 1566 (2018).

29. Konopleva, M., Contractor, R., Tsao, T., Samudio, I., Ruvolo, P., Kitada, S., Deng, X., Zhai, D., Shi, Y., Sneed, T., Verhaegen, M., Soengas, M., Ruvolo, V., McQueen, T., Schober, W., Watt, J., Jiffar, T.,… Andreeff, M. Mechanisms of apoptosis sensitivity and resistance to the BH3 mimetic ABT-737 in acute myeloid leukemia. Cancer Cell 10, 375–388 (2006).

30. Kotschy, A., Szlavik, Z., Murray, J., Davidson, J., Maragno, A., Le Toumelin-Braizat, G., Chanrion, M., Kelly, G., Gong, J., Moujalled, D., Bruno, A., Csekei, M., Paczal, A., Szabo, Z., Sipos, S., Radics, G., Proszenyak, A.,… Geneste, O. The MCL1 inhibitor S63845 is tolerable and effective in diverse cancer models. Nature 538, 477–482 (2016).

31. Hormi, M., Birsen, R., Belhadj, M., Huynh, T., Cantero Aguilar, L., Grignano, E., Haddaoui, L., Guillonneau, F., Mayeux, P., Hunault, M., Tamburini, J., Kosmider, O., Fontenay, M., Bouscary, D. & Chapuis, N. Pairing MCL-1 inhibition with venetoclax improves therapeutic efficiency of BH3-mimetics in AML. Eur. J. Haematol. 105, 588–596 (2020).

32. van Delft, M., Wei, A., Mason, K., Vandenberg, C., Chen, L., Czabotar, P., Willis, S., Scott, C., Day, C., Cory, S., Adams, J., Roberts, A. & Huang, D. The BH3 mimetic ABT-737 targets selective Bcl-2 proteins and efficiently induces apoptosis via Bak/Bax if Mcl-1 is neutralized. Cancer Cell 10, 389–399 (2006).

33. Birkinshaw, R., Gong, J., Luo, C., Lio, D., White, C., Anderson, M., Blombery, P., Lessene, G., Majewski, I., Thijssen, R., Roberts, A., Huang, D., Colman, P. & Czabotar, P. Structures of BCL-2 in complex with venetoclax reveal the molecular basis of resistance mutations. Nat. Commun. 10, (2019).

34. Tausch, E., Close, W., Dolnik, A., Bloehdorn, J., Chyla, B., Bullinger, L., Döhner, H., Mertens, D. & Stilgenbauer, S. Venetoclax resistance and acquired BCL2 mutations in chronic lymphocytic leukemia. Haematologica 104, E434–E437 (2019).

35. Tarantelli, C., Gaudio, E., Hillmann, P., Spriano, F., Sartori, G., Aresu, L., Cascione, L., Rageot, D., Kwee, I., Beaufils, F., Zucca, E., Stathis, A., Wymann, M. P., Cmiljanovic, V., Fabbro, D. & Bertoni, F. The Novel TORC1/2 Kinase Inhibitor PQR620 Has Anti-Tumor Activity in Lymphomas as a Single Agent and in Combination with Venetoclax. Cancers (Basel). 11, (2019).

36. Jiang, H., Lwin, T., Zhao, X., Ren, Y., Li, G., Moscinski, L., Shah, B. & Tao, J. Venetoclax as a single agent and in combination with PI3K-MTOR1/2 kinase inhibitors against ibrutinib sensitive and resistant mantle cell lymphoma (MCL). Br. J. Haematol. 184, 298 (2019).

37. Ricci, J. E. & Chiche, J. Metabolic reprogramming of non-Hodgkin’s B-cell lymphomas and potential therapeutic strategies. Front. Oncol. 8, 556 (2018).

38. Tailler, M., Lindqvist, L. M., Gibson, L. & Adams, J. M. By reducing global mRNA translation in several ways, 2-deoxyglucose lowers MCL-1 protein and sensitizes hemopoietic tumor cells to BH3 mimetic ABT737. Cell Death Differ. 26, 1766 (2019).

39. Leprivier, G. & Rotblat, B. How does mTOR sense glucose starvation? AMPK is the usual suspect. Cell Death Discov. 6, 27 (2020).

40. Almacellas, E., Pelletier, J., Manzano, A., Gentilella, A., Ambrosio, S., Mauvezin, C. & Tauler, A. Phosphofructokinases Axis Controls Glucose-Dependent mTORC1 Activation Driven by E2F1. iScience 20, 434–448 (2019).

41. De Bock, K., Georgiadou, M., Schoors, S., Kuchnio, A., Wong, B., Cantelmo, A., Quaegebeur, A., Ghesquière, B., Cauwenberghs, S., Eelen, G., Phng, L., Betz, I., Tembuyser, B., Brepoels, K., Welti, J., Geudens, I., Segura, I.,… Carmeliet, P. Role of PFKFB3-driven glycolysis in vessel sprouting. Cell 154, (2013).

42. Shi, L., Pan, H., Liu, Z., Xie, J. & Han, W. Roles of PFKFB3 in cancer. Signal Transduct. Target. Ther. 2017 21 2, 1–10 (2017).

43. Koppenol, W., Bounds, P. & Dang, C. Otto Warburg’s contributions to current concepts of cancer metabolism. Nat. Rev. Cancer 11, 325–337 (2011).

44. Vander Heiden, M., Cantley, L. & Thompson, C. Understanding the Warburg effect: the metabolic requirements of cell proliferation. Science 324, 1029–1033 (2009).

45. Abdel-Wahab, A. F., Mahmoud, W. & Al-Harizy, R. M. Targeting glucose metabolism to suppress cancer progression: prospective of anti-glycolytic cancer therapy. Pharmacol. Res. 150, (2019).

46. Bittremieux, M., Rovere, R. M. La, Akl, H., Martines, C., Welkenhuyzen, K., Dubron, K., Baes, M., Janssens, A., Vandenberghe, P., Laurenti, L., Rietdorf, K., Morciano, G., Pinton, P., Mikoshiba, K., Bootman, M. D., Efremov, D. G., Smedt, H. De,… Bultynck, G. Constitutive IP3 signaling underlies the sensitivity of B-cell cancers to the Bcl-2/IP3 receptor disruptor BIRD-2. Cell Death Differ. 26, 531 (2019).

47. Cerella, C., Muller, F., Gaigneaux, A., Radogna, F., Viry, E., Chateauvieux, S., Dicato, M. & Diederich, M. Early downregulation of Mcl-1 regulates apoptosis triggered by cardiac glycoside UNBS1450. Cell Death Dis. 6, (2015).

48. Monaco, G., Decrock, E., Akl, H., Ponsaerts, R., Vervliet, T., Luyten, T., De Maeyer, M., Missiaen, L., Distelhorst, C., De Smedt, H., Parys, J., Leybaert, L. & Bultynck, G. Selective regulation of IP3-receptor-mediated Ca2+ signaling and apoptosis by the BH4 domain of Bcl-2 versus Bcl-Xl. Cell Death Differ. 19, 295–309 (2012).

49. Decuypere, J.-P., Welkenhuyzen, K., Luyten, T., Ponsaerts, R., Dewaele, M., Molgó, J., Agostinis, P., Missiaen, L., Smedt, H. De, Parys, J. B. & Bultynck, G. Ins(1,4,5)P3 receptor-mediated Ca2+ signaling and autophagy induction are interrelated. Autophagy 7, 1472 (2011).

50. Cavalier, M. C., Kim, S. G., Neau, D. & Lee, Y. H. Molecular basis of the Fructose-2,6-bisphosphatase reaction of PFKFB3: Transition state and the C-terminal function. Proteins 80, 1143 (2012).

51. Jones, G., Willett, P., Glen, R. C., Leach, A. R. & Taylor, R. Development and validation of a genetic algorithm for flexible docking. J. Mol. Biol. 267, 727–748 (1997).

